# Optimal reliability, robustness and control of nucleus centering in fission yeast is contingent on nonequilibrium force patterning

**DOI:** 10.1101/2022.06.28.497980

**Authors:** Ishutesh Jain, Madan Rao, Phong T. Tran

## Abstract

Cells, such as fission yeast, center their division apparatus to ensure symmetric cell division, a challenging task when the governing dynamics is stochastic. Here we show that the spindle pole body (SPB) positioning, which defines the division septum, is controlled by the patterning of nonequilibrium polymerization forces of microtubule (MT) bundles. We define two cellular objectives, *reliability*, the mean SPB position relative to the geometric center, and *robustness*, the variance of the SPB position, which are sensitive to genetic perturbations that change cell length, MT-bundle number/orientation and MT dynamics. We show that the optimal control of reliability and robustness required to minimize septum positioning error is achieved by the wild-type (WT). A stochastic model for the MT-based nucleus centering, with parameters measured directly or estimated using Bayesian inference, recapitulates WT optimality. We use this to perform a sensitivity analysis of the parameters that control nuclear centering.

## 1 Introduction

The nucleus and other organelles in eukaryotic cells display precise and reproducible intracellular positioning (Chan & Marshall 2012, Marshall et al. 2012, Bornens 2008). For instance, the nucleus in sea urchin eggs and the mitotic spindle in *C. elegans*, localise at the center to within 5% of the cell length (Pécréaux et al. 2016, Minc et al. 2011). Reliable positioning is a consequence of an organised patterning of nonequilibrium intracellular forces that arise from structures that are cell spanning, adaptive, and simultaneously pliable and rigid, such as the active cytoskeleton (Jimenez et al. 2021).

Position control by fluctuating nonequilibrium forces from the active cytoskeleton appears to be ubiquitous across all eukaryotic life forms. This can arise as a result of *pulling forces* from actomyosin or motor-microtubule (MT) complexes or *pushing forces* from MT polymerization-depolymerization (Jimenez et al. 2021, Schuh 2011, Azoury et al. 2008, Almonacid et al. 2018, Colin et al. 2020, Makhija et al. 2015, Tran et al. 2001, Théry et al. 2007, Grill et al. 2003). For instance, nucleus positioning and spindle migration to the cell surface in mouse oocytes is driven by stresses exerted by actin cytoskeleton (Schuh 2011, Azoury et al. 2008, Almonacid et al. 2018, Colin et al. 2020, Makhija et al. 2015). Contractile stresses arising from actomyosin are also seen to be responsible for proper nuclear positioning in mouse fibroblast cells (Makhija et al. 2015, Rupprecht et al. 2018). In most cells, MT-cytoskeleton based active forces operate on the cortex either via pushing forces caused by MT polymerization against the cortex or pulling forces applied by anchored, minus-end directed motors or a combination of these (Tran et al. 2001, Théry et al. 2007, Grill et al. 2003).

Nonequilibrium stresses from both actomyosin contractility and polymerization-depolymerization of MT, exhibit strong mechanical fluctuations (Desai & Mitchison 1997, Howard & Hyman 2003, Mizuno et al. 2007, Gnesotto et al. 2018, Xiao et al. 2017). For instance, the growth dynamics of individual MT exhibits dynamical instability (Desai & Mitchison 1997, Howard & Hyman 2003, Mitchison & Kirschner 1984, Walker et al. 1988, Verde et al. 1992), with a well defined growth-phase before going to ‘catastrophe’ leading to a rapid shrinking-phase. Further, the number of MT involved in active force generation are subject to fluctuations, owing to the stochastic dynamics of nucleation and growth. Evidence suggests that the mechanical fluctuations of MTs are sensitive to applied forces (Dogterom & Yurke 1997, Janson et al. 2003, Laan et al. 2008); this implies that MT (as indeed actomyosin) are both active *force generators* and *force sensors*, aspects that are advantageous for adaptation and robust control.

In this paper we study the fidelity of nucleus centering in wild type (WT) and genetically perturbed fission yeast (*Schizosaccharomyces pombe*) cells, using a combination of live-cell microscopy and detailed statistical analysis. Fission yeast cells divide by medial fission and produce two daughter cells that are identical in size. Symmetric cell division is contingent on precise localisation of the division plane apparatus (Minc et al. 2011, Besson & Dumais 2011, Oliferenko et al. 2009, Barr & Gruneberg 2007, Fededa & Gerlich 2012, Rappaport 1986, Gundersen & Worman 2013, Hayles & Nurse 2001, Wühr et al. 2010, Tanimoto et al. 2016, 2018, Moorhouse & Burgess 2014). The position of the division-septum is determined by the centering of the nucleus at the onset of mitosis (Tolić-Nørrelykke et al. 2005, Daga & Chang 2005), which in turn is controlled by forces exerted by the dynamic microtubule cytoskeleton (Sawin & Tran 2006, Tran et al. 2001, Hayles & Nurse 2001). In the interphase cell, microtubules are organized in 3-4 polar bundles, in which the active plus-ends orient towards the cell tips, and are nucleated from the microtubule-organizing centers (MTOCs) immobilized on the nucleus. Microtubules growing from these bundles can reach the cell tips and exert pushing forces that apply compressive stresses. Consequently, the nucleus feels a force in a direction opposite to the microtubule growth during contact and moves in response. This exclusivity of the nuclear centering force agents, makes fission yeast an attractive model system to understand how the patterning and adaptation of the active force machinery influence the reliable and robust centering of the nucleus.

Using high-resolution imaging, we follow the dynamics of the spindle-pole-body (SPB), a proxy for nuclear movement, in various genetic backgrounds that alter cell length, number of MT bundles, MT organization, and MT growth dynamics. In each of these conditions, we measure the fidelity of centering in terms of *reliability* (*δ*) - the mean SPB position relative to the geometric center of the cell, and *robustness* (*σ*_*x*_, *σ*_*y*_) - the standard deviation of SPB positions in the longitudinal (long.) and transverse (trans.) directions. We show that the fidelity of nuclear centering directly affects the ability of the cell to place its septum accurately and divide at the center; the WT-condition achieves optimal *δ* and *σ*_*x*_ statistics, which manifests in low error in septum positioning. We identify the key control parameters associated with MT force-patterning and growth dynamics. In the process, we establish that for precise centering of the nucleus, apart from optimum MT number and orientation, the MT length scale (in the milieu of the yeast cell) needs to be matched with the cell length.

Next, we develop a stochastic model for the MT-based nucleus centering, with parameters measured directly from our experiments or estimated from our data using Bayesian inference (Gelman et al. 2013). The model recapitulates the optimality of the WT; using this, we perform a sensitivity analysis of the parameters that control nuclear centering.

Our model allows us to infer principles of robust and reliable nucleus positioning, leading to several immediate implications that we outline in the Discussion.

## 2 Results

### 2.1 Cell to cell variation in reliability and robustness of SPB positioning

Since the spindle-pole-body (SPB) localizes on the nucleus and is the nucleator of MT, it provides a convenient peg to follow the position of the nucleus. We monitor the dynamics of SPB both in wild-type (WT) and in genetic variants, using high spatio-temporal resolution live-cell imaging across 20-50 cells (see Fig. 1 and *Methods*). Simultaneously, we monitor aspects of the MT organization and dynamics that control the positioning of the nucleus. This includes the number of MT bundles, their orientation with respect to the long axis of the rod-shaped fission yeast cells, total MT mass and MT growth dynamics (Fig. 1j). The coordinate system used to measure the longitudinal and transverse displacements from the origin is described in Fig. 1c. To standardize the observations between WT and the various perturbations, we used a strain with the following fluorescent tags: Sid4-mCherry (SPB) and EnvyGFP-Atb2 (MT). This background enabled the long-term observation of SPB and MTs.

**Figure 1:**
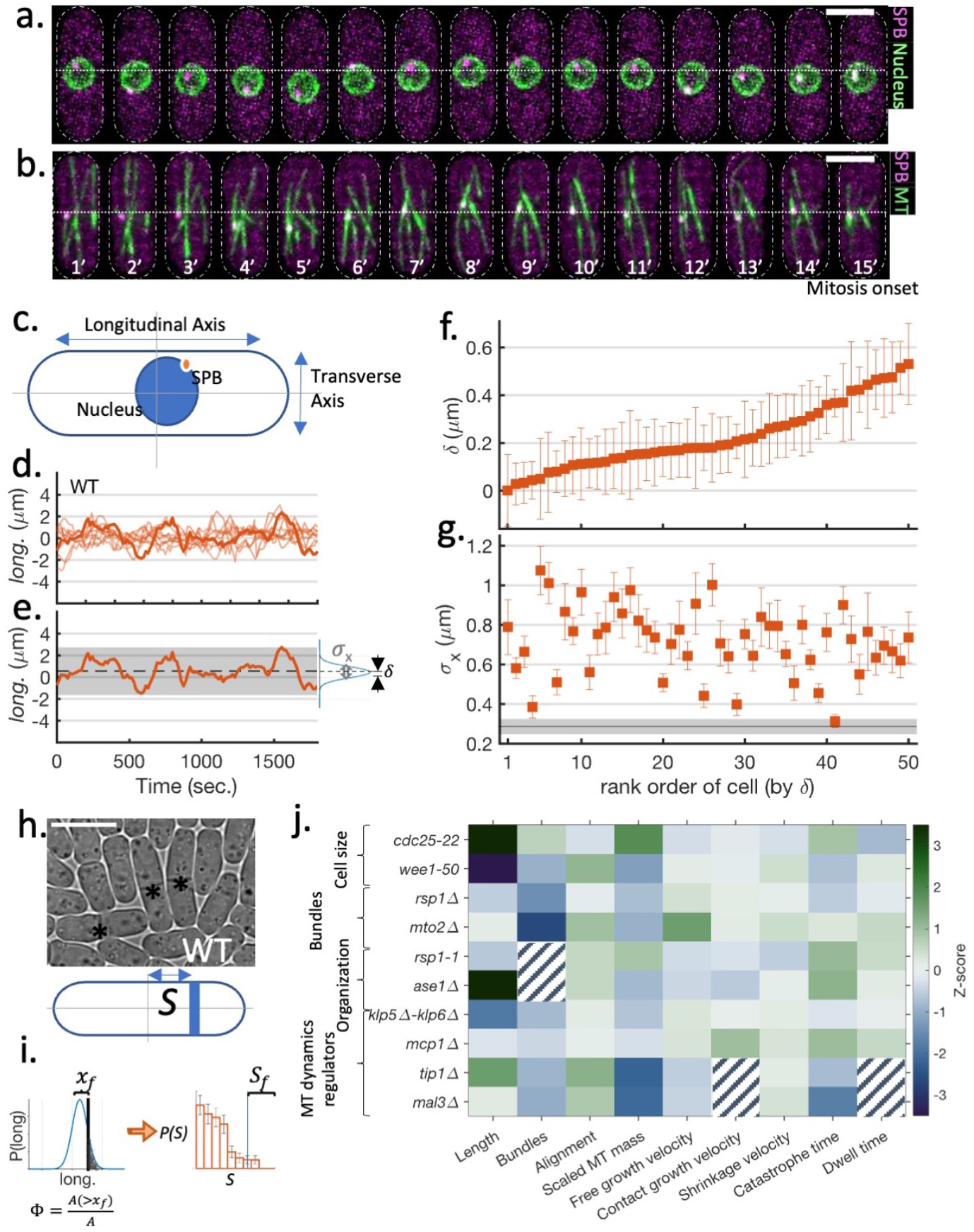
Dynamics of the nucleus and SPB in WT cells: **a.-b**. Time-lapse images of nucleus centering before the mitosis onset. (a) Representative example of the nuclear envelope (Cut11-GFP) and SPB (Sid4-mCherry) fluctuation in WT cell. (b) EnvyGFP-Atb2 Sid4-mCherry tagged strain (WT) showing the dynamics of SPB and MTs. Time-interval 1 min, scale-bar 10 *μm*. **c**. For each cell, we define an internal coordinate system with the origin at the cell centroid and the longitudinal axis aligned with the x-axis. By convention, the cells are oriented so that the time-average of longitudinal and transverse displacement of SPB w.r.t. the geometric center of the cell falls in the first quadrant. **d**. Example trajectories of longitudinal displacement of SPB. **e**. From such trajectories, we estimated the mean longitudinal displacement of SPB from the cell center (*δ*) and standard deviation in SPB position (*σ*_*x*_) for each cell. **f.-g**. Distribution of *δ* and *σ*_*x*_ in WT cells sorted by *δ*. The error bars show standard error, estimated using the moving-block bootstrap method with a block size equal to the autocorrelation time (Efron & Tibshirani 1994, Politis & Romano 1991). The black line (shaded area) in (g) is the mean *σ*_*x*_(±SE) in the MT destabilizing drug (MBC) treated cells. **h**. (Top) Examples of septum position in WT cells. (Bottom) The septum position ‘*S*’ is the distance of the septum from the cell center. **i**. To quantify the functional role of *δ* and *σ*_*x*_, we define a distance *x*_*f*_ from the cell center beyond which the septum localization is considered a *failure*. The fraction of cells (*S*_*f*_) whose septum localizes beyond *x*_*f*_, depends on *δ* and *σ*_*x*_. We calculate the *failure-coefficient* of SPB localization (Φ), by approximating the distribution of SPB as a Gaussian characterized by *δ* and *σ*_*x*_, as a readout to likely septum-mislocalization. Φ = *A*(> *x*_*f*_)/*A*, where 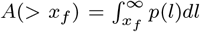 and 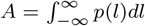. **j**. We use systematic perturbations in cell length, the number of MT bundles, MT organization, and MT growth dynamics regulator to understand the role of force patterning in determining SPB positioning. The heatmap shows the difference between various functional attributes between the WT and the mutant quantified using the 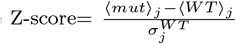, where ⟨…⟩ represent the mean, *j* is the feature index, and 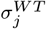 is the standard deviation observed in the WT-strain in the *j*^*th*^ feature (hatched squares denote cases where the features are ill defined).

We analyze the dynamics of the centroid of the nucleus and SPB in strains with GFP-Cut11 (nuclear envelop) and Sid4-mCherry (SPB) (see Fig. 1a and Fig. S1). Figure S1a shows the time-course of longitudinal displacement of the nucleus centroid and SPB for the WT-condition. We find that the longitudinal displacements of the SPB and the nucleus are strongly correlated (Pearson’s *r* = 0.58, *p* < 10^−57^), although the SPB experiences larger fluctuations than the nucleus. Treatment with MBC (Carbendazim), a MT-destabilizing drug, quenched these fluctuations (Fig. S1).

This suggests that the SPB-dynamics can be used as a proxy for nucleus localization before the onset of mitosis. Analysis of the time series of the longitudinal and transverse displacements, during a 30 min period, shows them to be statistically independent (Fig. S1). Moreover, the time-series is also stationary, with the mean and variance measured over a time window, being independent of time (Fig. S1). These measured quantities do not show any change as M-phase approaches, suggesting the dynamics is not subject to additional regulation. Consequently, the spatial position of SPB (and nucleus) at the onset of mitosis is causally determined by the dynamics prior to it.

From the time series of the longitudinal positions of the SPB in a given cell, we compute its stationary distribution, from which we extract *δ*, the mean SPB displacement from the geometric center, and *σ*_*x*_, the standard deviation of SPB displacement (Fig. 1e). *δ* is a measure of positioning *reliability*; smaller *δ* implies a more reliable centering. In ∼40% of WT cells, *δ* lies significantly away from the center, implying that even WT cells have a significant variation in reliability (Fig. 1f). On the other hand, *σ*_*x*_ is a measure of *robustness* in the positioning, and a smaller *σ*_*x*_ corresponds to a more robustly localized nucleus. In all WT cells, *σ*_*x*_ is ∼3-fold higher than the fluctuations observed after disintegrating MTs using MBC (Fig. 1g); thus, fluctuations arising from active forces makes a significant contribution to *σ*_*x*_. In principle, *δ* and *σ*_*x*_ represent two independent objectives for nucleus centering; changes in either lead to higher chances of an off-centered nucleus and, consequently, a high deviation in septum position (*S*) from the cell center (Fig. 1h-i).

In what follows, we will discuss changes upon specific genetic perturbations, keeping in mind that the perturbations inevitably change many attributes related to the organization and dynamics of MTs (quantified in Fig. 1j and Fig. S2-S6, and discussed further in detail) – we will take this into account in making our inferences.

### 2.2 SPB dynamics is sensitive to cell length

An aspect of cell geometry that is simple to vary is cell length. Since the MT-based pushing machinery that control nucleus positioning, span the entire length of the cell, perturbations of cell length can potentially inform about adaptability and scaling (Wühr et al. 2008, Hara & Kimura 2009, Hazel et al. 2013, Lacroix et al. 2018, Krüger et al. 2019).

We utilized the cell cycle mutant *cdc25-22* in which G2-M transition is delayed leading to longer cells; and *wee1-50* mutant strains, which shortens the cell cycle by inducing early G2-M transition, leading to shorter cells (Russell & Nurse 1986). Together with WT, these strains provide a 4x variation in cell length (Fig. 2a-c).

**Figure 2:**
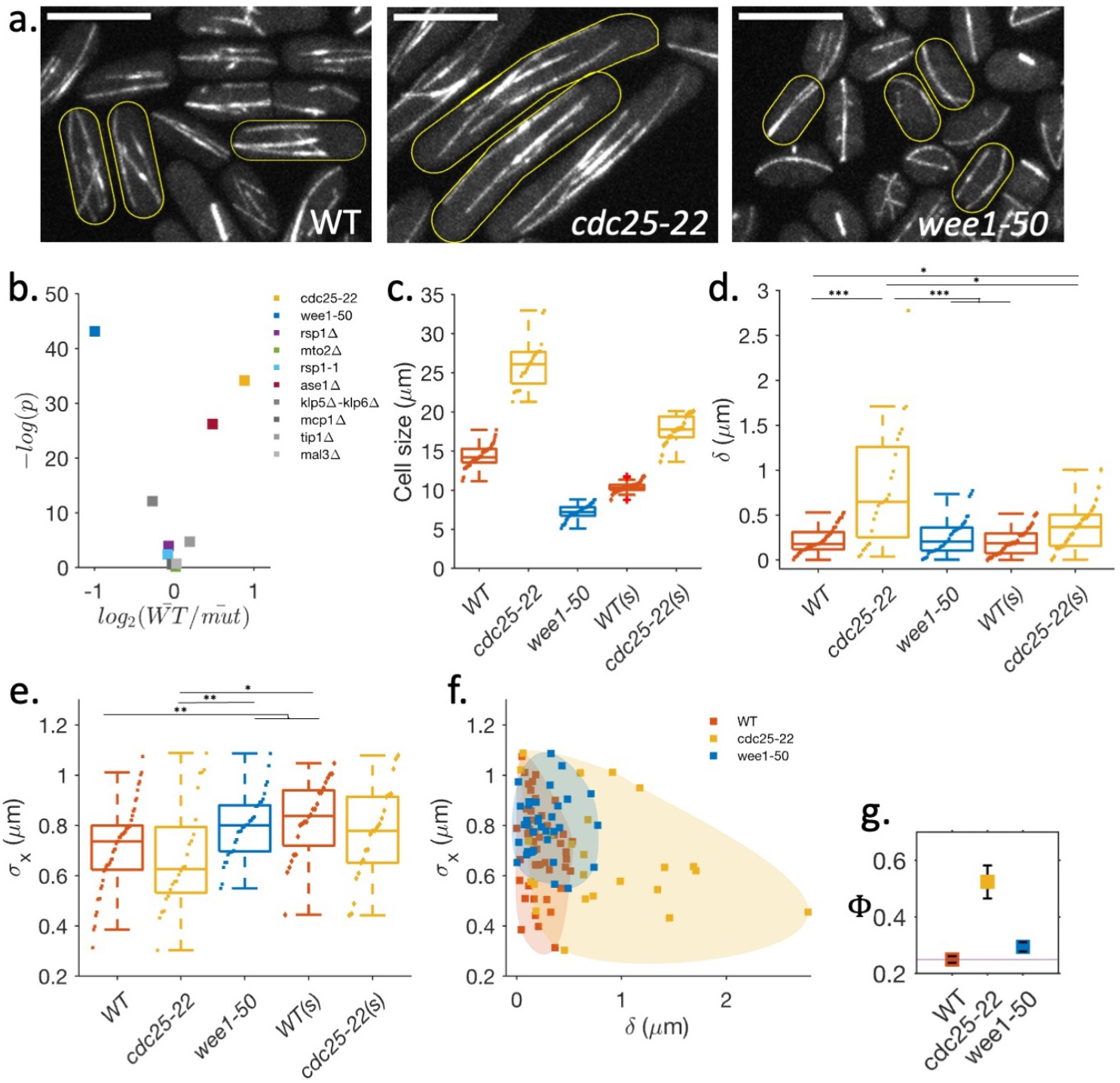
SPB-dynamics is sensitive to deviations of cell length from WT. **a**. MT organization in WT, *cdc25-22* (long), and *wee1-50* (small) cells. Scale bar 10 *μm*. **b**. Scatter plot of statistical significance (P-value) versus magnitude of change (fold change) in cell length w.r.t WT cells in studied strains. *cdc25-22*, and *wee1-50* show the most variation in the cell length. **c**. Cell length distribution at the onset of mitosis (for WT and *wee1-50*) and in the long *cdc25-22* cells (also Fig. S6). **d**. The mean longitudinal displacement of SPB from the cell center (*δ*) for each cell was obtained from SPB-dynamics data (see Fig. S2). *δ* increases in the long cells irrespective of the function of Cdc25. **e**. Standard deviation in SPB position (*σ*_*x*_) for each cell. *σ*_*x*_ increases in short cells. **f**. Scatter plot of *δ* and *σ*_*x*_ for WT, *cdc25-22, wee1-50* cells. A large fraction of *cdc25-22* have < larger than both WT and *wee1-50* and for a given value of *δ* small *wee1-50* have higher *σ*_*x*_ than the WT or *cdc25-22*. **g**. For each strain, we calculate the mean Φ as a function of *δ* and *σ*_*x*_ as a measure of nucleus mislocalization. (error bars are SE). [*** for *p* ≤ 10^−4^, ** for *p* ≤ 0.005, * for *p* < 0.05, Wilcoxon rank sum-test.]

The basic organization of MT-cytoskeleton in these strains is similar - MTs are arranged in bundles, mostly aligned with the long axis of the cell (Fig. 2a, Fig. S2). However, in the short *wee1-50* cells, MTs are more bent, resulting in a dispersed local MT orientation (Fig. S2). The number of MT-bundles and the net MT mass per unit length is large in *cdc25-22* cells and small in *wee1-50* cells compared to the WT, suggesting a systematic variation with cell length (Fig. S2). Importantly, measurement of the MT growth parameters (see Fig. S7), reveal that the free growth velocity, shrinkage velocity, and growth velocity during contact of MT to cell cortex are very similar to the WT (Fig. S2). The catastrophe time, however, shows a slight variation, averaging 115s, 148s, and 90s for the WT, *cdc25-22*, and *wee1-50* cells, respectively (Fig. S2). Similarly, the mean dwell times are 45 s for the WT, 17 s for *cdc25-22*, and cells 54s for *wee1-50*. Taken together, these results suggest that cell length perturbations have only a moderate effect on the basic MT-organization and intrinsic growth parameters. However, the dynamic parameters that rely on the interaction between MT and cell-cortex, e.g. the catastrophe and dwell times (Janson et al. 2003), and MT-mass show a dependence on cell length.

We now monitor the SPB-dynamics before the onset of mitosis both in the WT and these strains. A visual inspection of the time series of the longitudinal positions, immediately reveals that *wee1-50* cells exhibit larger fluctuations than the WT cells, while the *cdc25-22* cells show smaller fluctuations than the WT (Fig. S2). Additionally, in many *cdc25-22* cells, the mean SPB localization is visibly off-center. Consequently, many cells in the *cdc25-22* strain have large values of *δ* compared to the WT or *wee1-50* cells, suggesting inadequate reliability of positioning. This difference is independent of Cdc25 function, since in small *cdc25-22* cells, *δ* decreases (Fig. 2d). We see that *wee1-50* cells have significantly large *σ*_*x*_ in comparison to WT or *cdc25-22* cells (Fig. 2e), suggesting a less robust localization of nucleus in small cells. This difference is independent of Wee1 function since small WT cells also have large *σ*_*x*_ (Fig. S2). We conclude that an increase (decrease) in cell length from the typical WT, leads to less reliable (less robust) nucleus centering, respectively.

The observations of high *σ*_*x*_ in the shorter cells is reminiscent of the high fluctuations seen in *in vitro* assays where MT lengths are much longer than the compartment size (Laan et al. 2012, Faivre-Moskalenko & Dogterom 2002). The buckled MTs appear less dynamic and are associated with lower *σ*_*x*_ (Howard 2006).

Together, reliability (*δ*) and robustness (*σ*_*x*_) are the two independent phenotypic objectives for nucleus position at the onset of mitosis; good performance in these two objectives correspond to low values of both *δ* and *σ*_*x*_. We can ask how each cell in these strains fare in the realisation of these joint objectives. Owing to underlying cell-to-cell variations, we expect a cloud of points to represent each strain (the color-shaded region in Fig. 2f) in an objective space. Large cdc25-22 cells occupy an expanded space where many cells have larger *δ* than the WT and wee1-50 cells. Conversely, small wee1-50 cells occupy a region where many cells have higher *σ*_*x*_ than the WT cells. These observations suggest that collective minimization of *δ* and *σ*_*x*_ is sensitive to the cell length.

An increase in either *δ* or *σ*_*x*_ enhances the chances of mislocalization of the septum. We define the *failure-coefficient* of SPB localization (Φ), that quantifies the likelihood of extreme septum localization events (Fig. 1i) beyond a fixed threshold, say the extreme-5% septum localization events in the WT cells. Comparing Φ across the three strains suggests that the chance of mislocalization is the least in the WT cell length (Fig 2g).

### 2.3 Robustness in nucleus positioning increases with number of MT-bundles

We now look at how one may systematically vary parameters characterising MT-based force patterning. Here, we study how variation in MT-bundle numbers affects nucleus positioning. WT cells have 3-4 MT bundles which consist of upto 12 filaments arranged in a predominantly anti-parallel manner (Sawin & Tran 2006, Janson et al. 2007, Höög et al. 2007, Carazo-Salas & Nurse 2006). We study deletion mutants where this bundle number can vary - thus *rsp1*Δ have 2-3 and *mto2*Δ have 1-2 MT bundles (Fig. 3a-c). Rsp1 localizes to interphase-MTOCs (iMTOCs), non-SPB MTOCs which are only present during interpahse, and is required for the proper functioning of iMTOCs (Zimmerman et al. 2004, Shen et al. 2018). Mto2 is a component of the *γ*-tubulin activator complex, and deletion of Mto2 leads to nonfunctioning iMTOCs, which results in SPB being the only active MTOC (Janson et al. 2005, Sawin & Tran 2006, Lynch et al. 2014). Both these mutants have cell-size similar to WT-cells at the onset of mitosis (Fig. S3).

**Figure 3:**
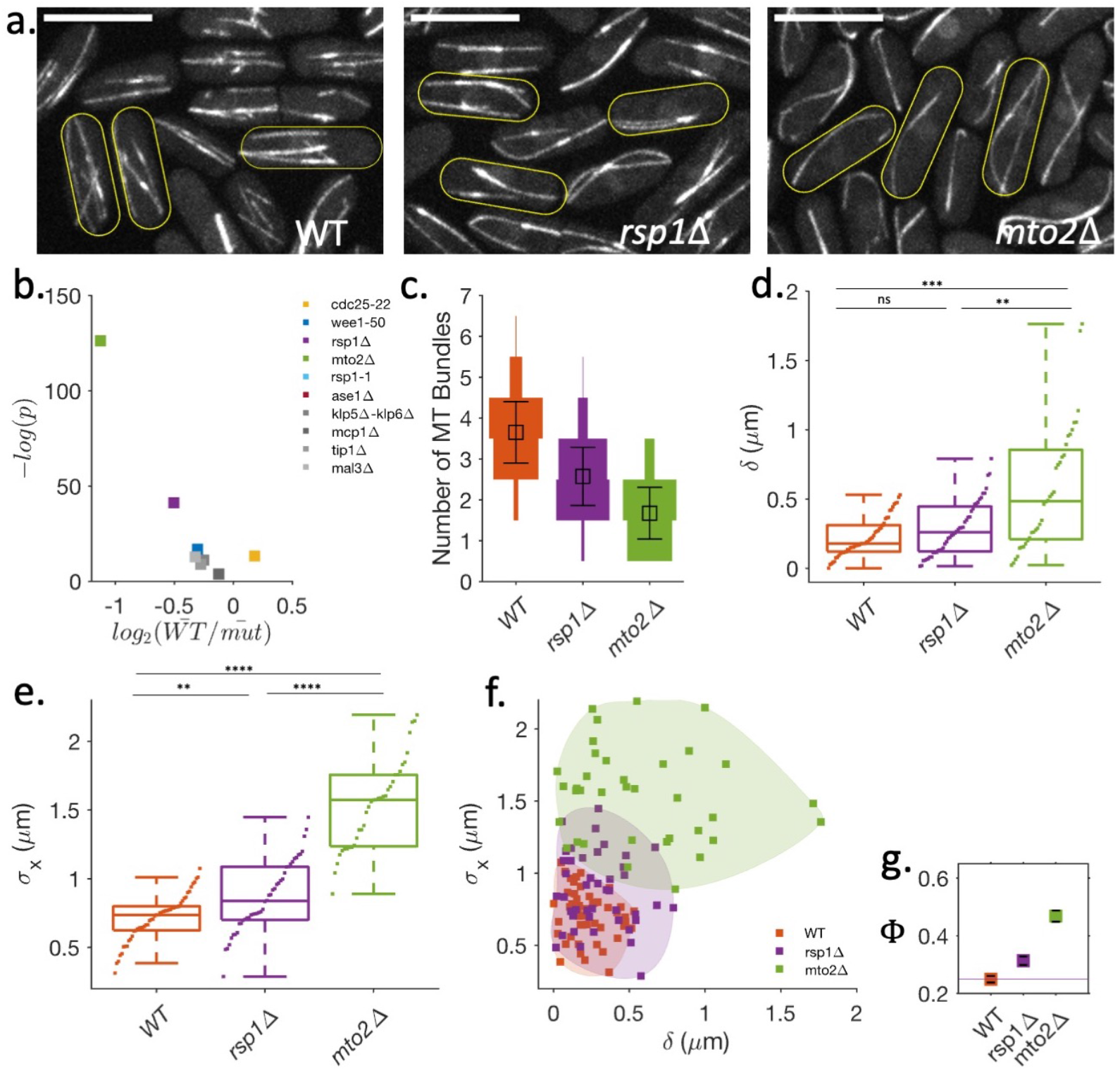
Fidelity of nucleus centering is affected by the number of MT-bundles. **a**. MT organization in WT, *rsp1*Δ, and *mto2*Δ cells. Scale bar=10 *μm*. **b**. Scatter plot of statistical significance (P-value) versus magnitude of change (fold change) in the number of MT bundles w.r.t WT cells in studied strains. **c**. Distribution of the number of MT bundles with a mean equal to 3.7 (WT), 2.5 (*rsp1*Δ), and 1.6 (*mto2*Δ) (error bars are SD) (also Fig. S6). **d**. Mean longitudinal displacement of SPB from the cell center (*δ*) for each cell obtained from SPB-dynamics data (see Fig. S3). **e**. Standard deviation in SPB position (*σ*_*x*_) for each cell. *σ*_*x*_ increases with a decrease in the number of MT-bundles. **f**. Scatter plot of *σ*_*x*_ and *δ* for WT, *rsp1*Δ, and *mto2*Δ cells. For a given *δ, σ*_*x*_ follows the trend WT<*rsp1*Δ<*mto2*Δ. **g**. The mean Φ as a function of *δ* and *σ*_*x*_ as a measure of nucleus mislocalization. (error bars are SE). [** ** for *p* ≤ 10^−5^, *** for *p* ≤ 10^−4^, ** for p ≤ 0.005, * for *p* < 0.05, ns is not significant. Wilcoxon rank-sum test.]

The *rsp1*Δ and *mto2*Δ cells have very similar MT-mass, though significantly reduced in comparison with WT (Fig. S3). Not surprisingly, *mto2*Δ cells show a higher fraction of bent MTs (Fig. S3). We find that most of the dynamic parameters associated with MT-growth are not significantly different from the WT, except for the free MT-growth velocities which are elevated in the *mto2*Δ strain (Fig. S3; Table S1). High MT-growth velocities and similar MT-catastrophe times are consistent with the observation that *mto2*Δ cells have longer MTs than WT and *rsp1*Δ cells. Further, having the same MT-mass for longer MT suggests that *mto2*Δ has fewer MTs numbers than *rsp1*Δ cells. In conclusion, changes in MT-bundles in these strains also changes the MT-numbers and one should expect the number of MTs to follow the order WT>*rsp1*Δ>*mto2*Δ.

Next, we analyzed the SPB dynamics in these strains 30 mins before the onset of mitosis (Fig. S3). We note that in some *mto2*Δ cells, the SPB movement ceases much before the onset of mitosis. This was observed earlier (Janson et al. 2005) and attributed to an SPB-MT bundle detachment phenotype, we therefore exclude these cells from our analysis. As seen in Fig. S3f, the SPB in *rsp1*Δ and *mto2*Δ cells appear much more dynamic than in WT cells. We quantified the reliability *δ* and robustness *σ*_*x*_ of nucleus positioning for each cell (Fig. 3d, e). The values of *δ* are significantly higher in *mto2*Δ cells compared to WT and *rsp1*Δ cells, suggesting that many *mto2*Δ cells have less reliable positioning of the nucleus. However, *σ*_*x*_ appears to increase with a decrease in the number of MT bundles i.e., WT<*rsp1*Δ<*mto2*Δ. This relationship becomes more evident in the objective space of *δ* and *σ*_*x*_ (Fig. 3f). Moreover, the transverse fluctuations of the SPB are higher in *mto2*Δ than in WT (see Fig. S3). This suggests that having multiple-bundles also aids in stabilizing the transverse motion of the nucleus.

In comparison to WT-strain, *rsp1*Δ and *mto2*Δ strains show a much larger cell-to-cell variation in objective space (Fig. 3f). We estimate the failure-coefficient of SPB localization, Φ, using the values of *δ* and *σ*_*x*_ for each condition, and find that it depends on the number of MT bundles. We find that the WT with the larger number of MT bundles, has the least Φ. There is no appreciable gain beyond a certain number of MT bundles, suggesting a tradeoff between effectiveness and economy in cellular costs.

### 2.4 Variation in orientational patterning of MT filaments and bundles

As stated, the MT in WT cells is organized as linear bundles aligned along the long axis of the cell. In contrast, *rsp1-1* mutant cells have MTs that have an aster-like organization, and MT-crosslinker, Ase1, deletion strain (*ase1*Δ) have disorganized and non-bundled MTs (Fig. 4a) (Zimmerman et al. 2004, Loiodice et al. 2005). While *rsp1-1* cells have similar cell length as WT, in our experiments, the typical length of *ase1*Δ cells is longer than WT at the onset of mitosis; therefore we also analyze small *ase1*Δ cells (*ase1*Δ(s)) (Fig. S4). The mean scaled MT-mass in *rsp1-1* (*ase1*Δ) cells is slightly higher (lower) than in WT cells (Fig. S4). The orientation distribution of MT filaments is much broader in *rsp1-1* and *ase1*Δ cells (Fig. 4b, c) than in WT, with many filaments showing large angular deviations from the long axis.

**Figure 4:**
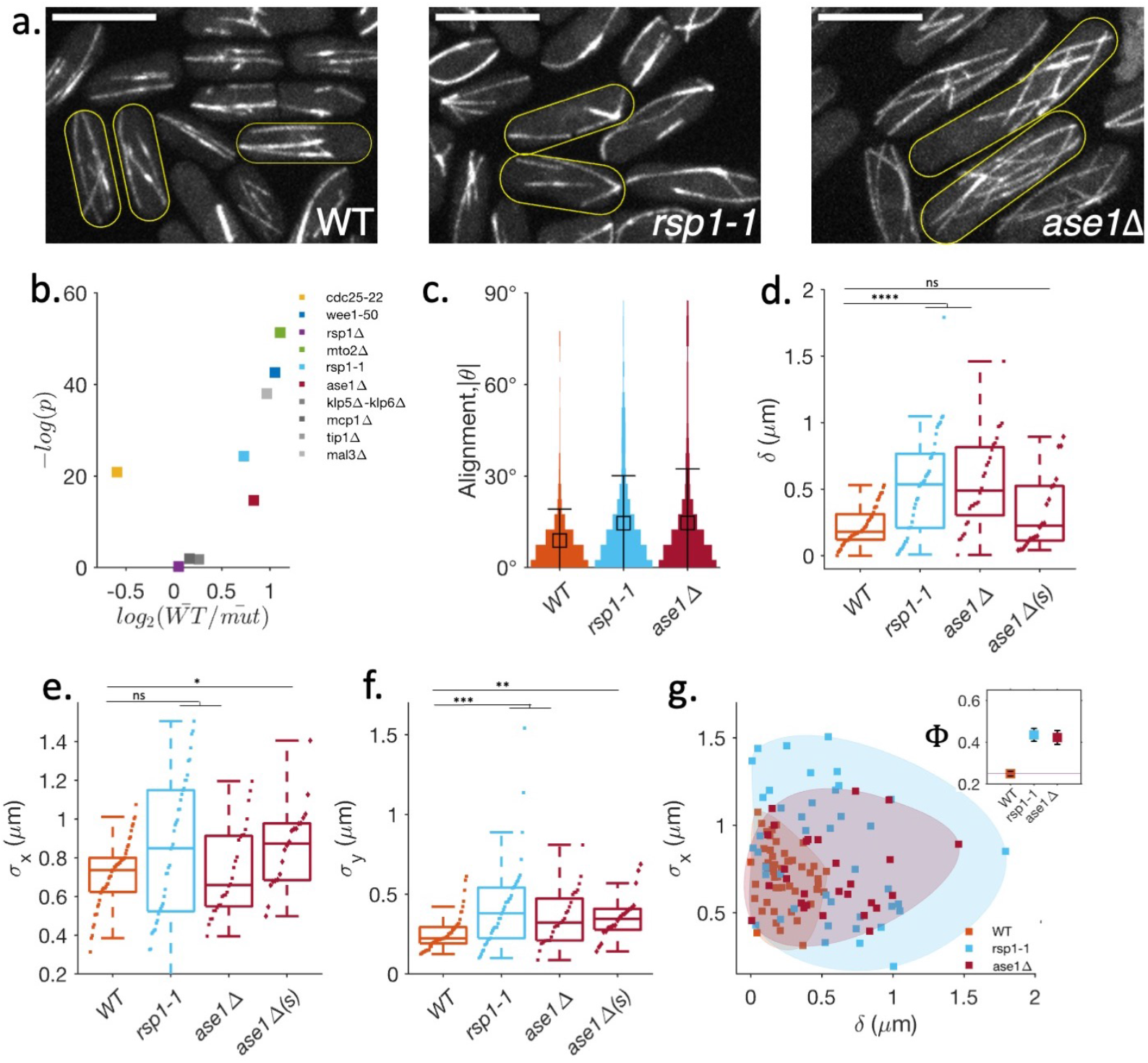
Orientation patterning of MTs affects the fidelity of nucleus centering. **a**. MT organization in WT, *rsp1-1*, and *ase1*Δ cells. In *rsp1-1* cells, MTs are arranged in an aster-like fashion, whereas in *ase1* Δ cells, MTs are unbundled and disorganized. Scale bar = 10 *μm*. **b**. Scatter plot of statistical significance (P-value) versus magnitude of change (fold change) in the alignment of MT w.r.t WT cells in studied strains. Apart from *rsp1-1* and *ase1*Δ, *wee1-50, mto2*Δ, and *mal3*Δ also have a large deviation in MT alignment. However, in *wee1-50* and *mto2*Δ, this large deviation is partly a result of buckled MTs; in the *mal3*Δ cells mean MT length is small. Moreover, *rsp1-1* and *ase1*Δ have distinct unbundled MT organization. **c**. Distribution of alignment of MTs quantified by the absolute local angle |*θ*| of MT filament relative to the longitudinal axis of the cell. Both *rsp1-1* and *ase1*Δ cells show large angular deviation. **d**. Mean longitudinal displacement of SPB from the cell center (*δ*) for each cell obtained from SPB-dynamics data (see Fig. S4). **e**. Standard deviation in SPB position (*σ*_*x*_) for each cell. **f**. Transverse fluctuations of SPB (*σ*_*y*_) in WT, *rsp1-1, ase1*Δ and small *ase1*Δ cells. **g**. Scatter plot of *δ* and *σ*_*x*_ for WT, *rsp1-1* and *ase1*Δ cells. Both *rsp1-1* and *ase1*Δ occupy a wider region in the objective space. Inset: The mean Φ as a function of *δ* and *σ*_*x*_ as a measure of nucleus mislocalization. (error bars are SE). [** ** for *p* ≤ 10^−5^, ** * for *p* ≤ 10^−4^, ** for p ≤ 0.005, * for *p* < 0.05, ns is not significant. Wilcoxon rank-sum test.]

Despite this, most MT-growth parameters in these strains are very similar to the WT (Fig. S4). The only significant difference is in catastrophe time, which is higher in both *rsp1-1* (154 s) and *ase1*Δ (157 s) strains in comparison to WT (115 s) (see Table S1 in SI).

Fig. S4e shows typical SPB-trajectories in these cells during the 30 min interval before the onset of mitosis. In some *rsp1-1* cells, the SPB pauses for an extended period near the pole. These typically happen when the MTs are in an aster-like organisation.

In *ase1*Δ cells, the mean position of the SPB is displaced away from the cell center. Analysis of *δ* and *σ*_*x*_ for *rsp1-1* and *ase1*Δ cells (Fig. 4d, e), shows that *δ* is more widely distributed, and that the mean *δ* is significantly higher than in the WT. On the other hand, *σ*_*x*_ in *rsp1-1* and *ase1*Δ cells are not significantly different from WT cells. Since *ase1*Δ cells are typically longer than the WT cells, we also study the dynamics of SPB-positioning in small *ase1*Δ cells (marked *ase1*Δ(s), also see Fig. S4 in SI). In small *ase1*Δ cells, while the values of *δ* are similar to WT cells, *σ*_*x*_ is higher. Moreover, *rsp1-1* and *ase1*Δ cells displayed larger fluctuations in the transverse displacements of SPB (Fig. 4f).

In the objective space of *δ* and *σ*_*x*_, both *rsp1-1* and *ase1*Δ strain show large cell-to-cell variation, and the failure-coefficient of SPB localization (Φ) increases in both of these strain (Fig. 4g). Thus, having linear bundles of MT aligned along the long axis (as in WT) leads to a more reliable and robust nucleus centering.

### 2.5 Optimal centering of nucleus requires favourable MT growth dynamics

Finally, we systematically studied how variation in the MT-growth dynamics affects nucleus centering using the mutants, (i) a deletion strain of Mal3 (*mal3*Δ), an EB1 homolog; (ii) a deletion strain of Tip1 (*tip1*Δ), a CLIP-170 homolog (both are MT-growth factors); (iii) a deletion strain of Klp5 and Klp6 (*klp5*Δ-*klp6*Δ), kinesin-8 proteins, and (iv) an Mcp1 deletion strain (*mcp1*Δ), an MT-binding protein (both are MT-catastrophe factors) (Brunner & Nurse 2000, Zheng et al. 2014, Meadows et al. 2018, Emanuel et al. 2004, Tischer et al. 2009) (Fig. 5a and Fig. S5). We find that *klp5*Δ-*klp6*Δ cells are shorter and *tip1*Δ cells are longer at mitosis compared to WT (Fig. S5). While the MT-mass relative to cell length is reduced in *tip1*Δ and *mal3*Δ strains as expected (Brunner & Nurse 2000, Howard & Hyman 2003, Akhmanova & Steinmetz 2008), we also observe a significant reduction in MT-mass in *klp5*Δ-*klp6*Δ strain. This may reflect a decrease in the nucleation activity consistent with the *in vitro* observation that Klp5-Klp6 acts as a nucleation factor (Erent et al. 2012) (Fig. 5b, c). MT orientation in *klp5*Δ-*klp6*Δ and *mcp1*Δ cells are similar to the WT-cells (Fig. S5). However, in *tip1*Δ and *mal3*Δ cells, the MTs are less aligned along the cell-axis, highlighting the significance of long MTs (Hayles & Nurse 2001, Deinum & Mulder 2013).

**Figure 5:**
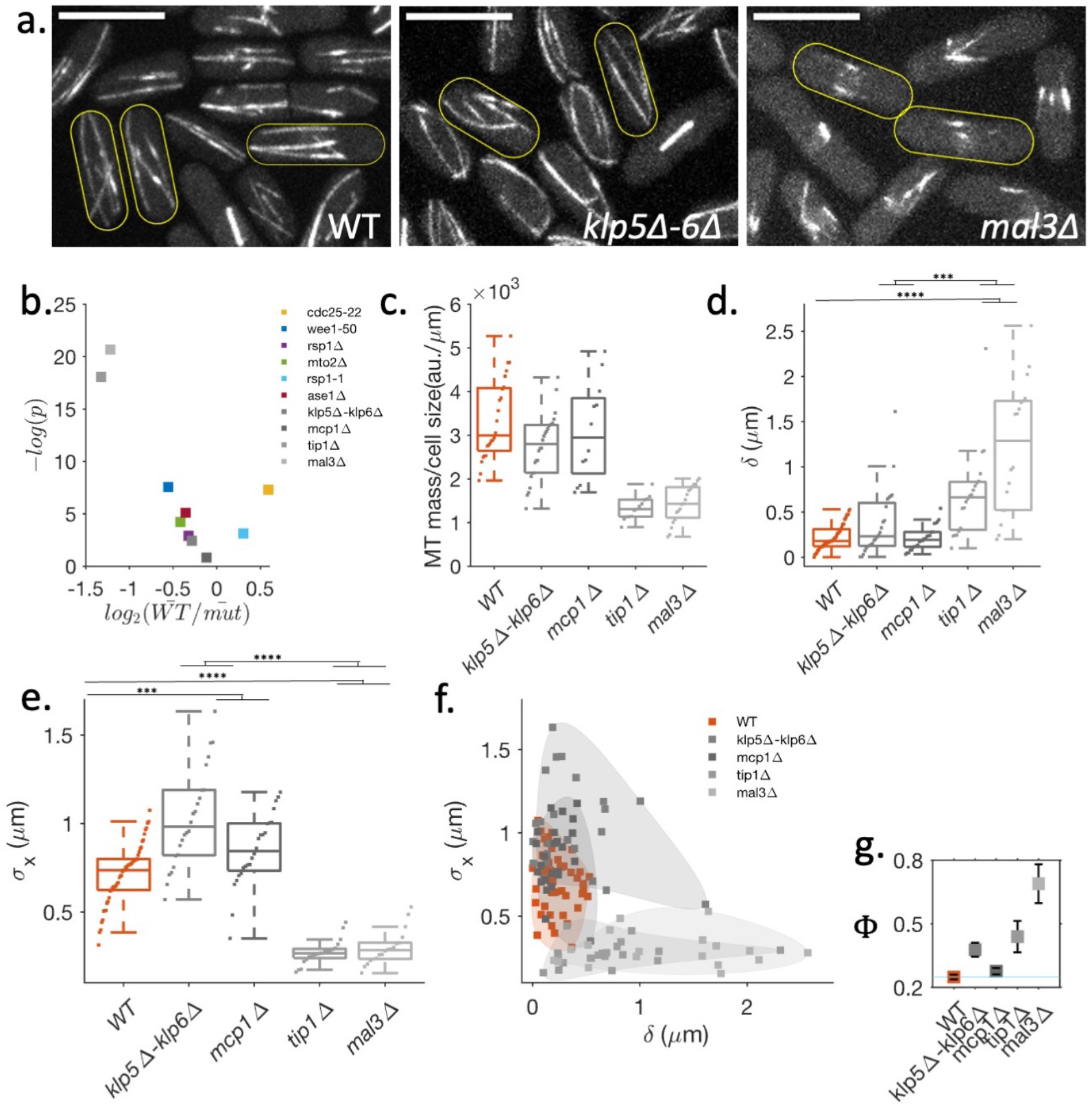
Optimal centering of the nucleus requires favorable MT growth dynamics. **a**. MT organization in MT-growth dynamics impaired mutants (also Fig. S5). Scale bar 10 *μm*. **b**. Scatter plot of statistical significance (P-value) versus magnitude of change (fold change) in the scaled MT-mass w.r.t WT cells in studied strains. *mal3*Δ and *tip1*Δ (both are MT growth factors) have low MT mass. However, we also observed a reduction in MT-mass in *klp5*Δ-*klp6*Δ. **c**. MT-mass, measured from the intensity of MT fluorescence, scaled with cell length (also Fig. S6). **d**. Mean longitudinal displacement of SPB from the cell center (*δ*) for each cell obtained from SPB-dynamics data (see Fig. S5). **e**. Standard deviation in SPB position from the cell center (*σ*_*x*_) for each cell. **f**. Scatter plot of *δ* and *σ*_*x*_. The *mal3*Δ and *tip1*Δ strains, with attenuated MT-growth, have large *δ* in the majority of the cells. In *klp5*Δ-*klp6*Δ and *mcp1*Δ strains, many cells have much higher *σ*_*x*_ than the WT. **g**. The mean Φ as a function of *δ* and *σ*_*x*_ as a measure of nucleus mislocalization. (error bars are SE). [** ** for *p* ≤ 10^−5^, *** for *p* ≤ 10^−4^, ** for *p* ≤ 0.005, * for *p* < 0.05, ns is not significant. Wilcoxon rank-sum test.]

The MT-growth dynamics in these strains have the following features (Fig. S5): Both MT-growth and shrinkage velocities are very similar in these strains. The growth-velocities of MT during contact with cell-tip are also similar in WT, *klp5*Δ-*klp6*Δ and, *mcp1*Δ cells. However, MT undergoes catastrophe much before reaching the cell-tip in *mal3*Δ and *tip1*Δ cells, consequently showing higher catastrophe frequency. In our experimental sampling, we do not see much difference between MT-catastrophe frequency in WT and *klp5*Δ-*klp6*Δ and *mcp1*Δ cells. However, for *mcp1*Δ cells, we do notice a wider distribution of both MT-catastrophe time and dwell times.

Next, we characterized the reliability and robustness of the nucleus centering in these cells (Fig. 5d, e). Both, *klp5*Δ-*klp6*Δ and *mcp1*Δ cells show higher fluctuations in SPB-dynamics compared to WT. In contrast, *tip1*Δ and *mal3*Δ cells, in which the MT-catastrophe rate is higher thus leading to shorter MTs, show suppressed SPB-dynamics away from the cell center (Fig. S5). This leads to high value of *δ* (Fig. 5d), similar to the long *cdc25-22* cells. However, both *klp5*Δ-*klp6*Δ and *mcp1*Δ cells show much higher *σ*_*x*_ (Fig. S5 and Fig. 5e). Earlier, high *σ*_*x*_ in *klp5*Δ-*klp6*Δ has been attributed to the lack of enhanced catastrophe at cell ends (Glunčić et al. 2015), however, we propose that a reduction in MT might also contribute to this phenomenon. Cumulative analysis of *δ* and *σ*_*x*_ in the objective space (Fig. 5f) shows that *mal3*Δ and *tip1*Δ prominently occupy a space to the right of the WT with smaller *σ*_*x*_, and *klp5*Δ-*klp6*Δ and *mcp1*Δ cells occupy space above the WT with small *δ*. The failure-coefficient Φ of SPB localization (Fig. 5g) shows that the altered MT-growth dynamics increase the chance of nucleus mispositioning.

### 2.6 Precise nucleus centering is key to medial positioning of division plane

A crucial function of nuclear positioning in fission yeast at the onset of mitosis, is to define the site of septum formation. We have shown that cell length, MT-bundle numbers, MT-architecture and, MT growth dynamics influence both reliability and robustness of nucleus centering. We next ask how do these traits impact the septum position. To this end, we observed the septum position in the studied strains at 35°C. (Note that *cdc25-22* cells do not commit to mitosis at 35°C. However, we found that *cdc25-22* cells growing at 35°C immediately go to mitosis on changing the temperature to 25°C (see *Methods*). The off-center septum is visible in many strains. We quantify the error in septum positioning by measuring the distance of the septum from the cell center (*S*) (Fig. 1h). Fig. 6b show the cumulative distribution of *S* in these strains. It is immediately clear that all mutants show a much broader distribution of septum positions than WT. Both, high *δ* or high *σ*_*x*_ of the SPB-position get mapped onto a broad distribution of *S*. For example, the *wee1-50* and *mcp1*Δ strains have higher *σ*_*x*_ than WT, and *ase1*Δ have higher *δ* than WT, even though the *σ*_*x*_ is comparable - all these strains exhibit a much broader *S* distribution.

**Figure 6:**
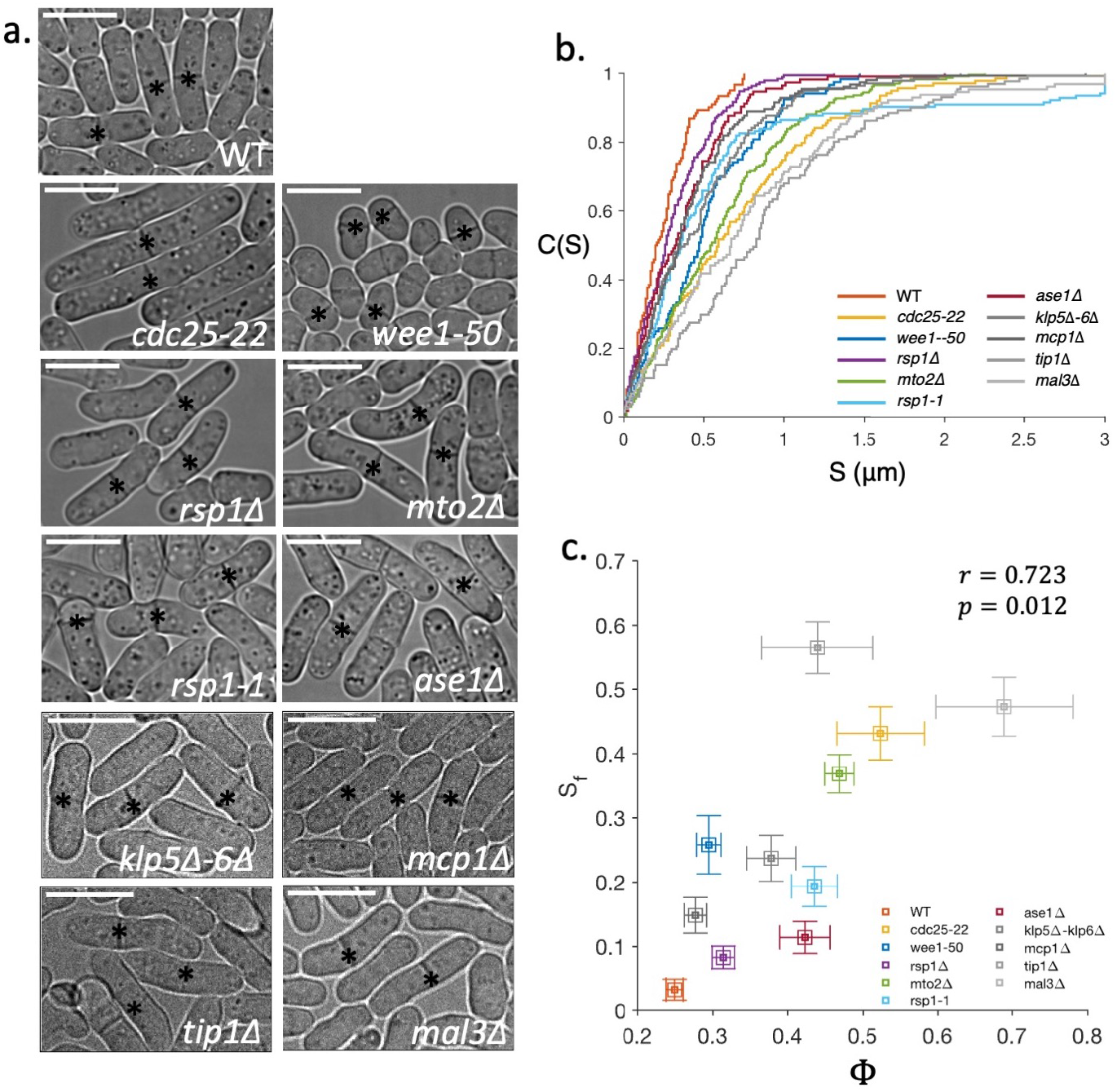
Proper positioning of the septum is contingent on nucleus centering. **a**. Panel shows examples of septum location in different strains. Scale bars 10 *µm*. Asterisks (∗) mark the septa. **b**. Cumulative distributions of S for various stains (also Fig. S6 for p-values between these strains). **c**. Fraction of cells with failed septum localization (*S*_*f*_) correlates with mean failure-coefficient of SPB localization Φ for the different strains, indicating the relevance of the combined optimization of *δ* and *σ*_*x*_. Error bars represent standard error. The error bar on *S*_*f*_ is estimated using 100 boot-strapped replicas.

Based on these observations, we expect that *S*_*f*_, the failed septum localization (fraction *S* that lies beyond *x*_*f*_), should be strongly correlated with Φ (fraction of cells with failed septum localization), as Φ is a function of both *δ* and *σ*_*x*_ (Fig. 6c). Pearson correlation statistics between the mean Φ and *S*_*f*_ show a correlation of 0.72 (*p* < 0.05). We note that some of the mutants explored here are also known to show some degree of morphological variations; e.g., *tip1*Δ, *mal3*Δ have noticeably bent cells in their population. These variations have been linked to the role MTs plays in the disposition of polarity factors (Hayles & Nurse 2001, Martin 2009, Minc et al. 2009). Given these complexities, we contend that the ∼ 72% variation in *S*_*f*_ explained by Φ reflects the high-influence of optimal nucleus centering on the septum position.

### 2.7 Stochastic model of MT-driven nucleus centering recapitulates optimality

We next develop a stochastic model to explain how the underlying MT force patterning may lead to an optimum nucleus centering. We have seen that the position of the nucleus in the interphase cell is controlled by forces exerted by the dynamic MTs which nucleate at the nucleus and are organized in a finite number of bundles. MTs polymerizing from these bundles can reach the cell tips and exert pushing forces, undergo dynamical instability and shrink. Consequently, the nucleus feels a stochastic force in a direction opposite to the MT growth during the contact phase and moves in response.

This is summarized in the following dynamical equations in terms of nonequilibrium forces and torques on the nucleus:

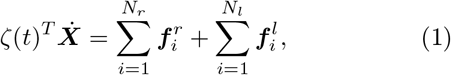

and

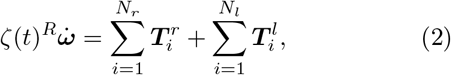

where ***X*** and ***ω*** are the position of the centroid of the nucleus (taken as a rigid sphere) and angle between the longitudinal axis of the cell and the vector connecting the centroid of the nucleus to SPB, respectively. The effective translational (*T*) and rotational (*R*) drag matrices *ς*(*t*)^*T*/*R*^, are a combination of the drag coefficients of the nucleus and MT-bundles. The force ***f***^*r*/*l*^ and the torque ***T***^*r*/*l*^ are applied by *N*^*r*/*l*^ MTs directed towards the right (*r*) and left (*l*) cell-tips (see detailed description in SI section A.2). The nucleus experiences a pushing force only when the MTs are in contact with the right or left cell boundary,

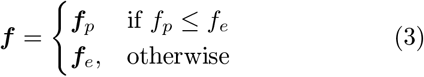

where ***f***_*p*_ is the MT-polymerization driven pushing force described by the force-velocity relationship: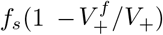, where *f*_*s*_ is the stall force and *V*_+_ and 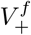 is the growth velocity of MT during the free growth and dwell phase, and ***f***_*e*_ is the critical Euler buckling force of MTs, which depends on the MTs flexural rigidity *k* (Table S2 in SI). Contact with the cell tips lasts only till the onset of catastrophe, wherein the MTs shrink rapidly and exert zero force till it recovers, grows and reaches the cell tip again. The catastrophe times *τ*_*cat*_ and their recovery are stochastic, and we measure their statistical distribution. In our analysis, we include a force dependent reduction in catastrophe time via the relation 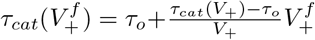, where *τ*_*o*_ is the mean catastrophe time at *V*_+_ = 0 (Janson et al. 2003, Foethke et al. 2009). We measure these growth (and shrinkage) velocities in the WT and the different mutant strains, and find them similar (Fig. S2-S5, Table S1 in SI).

We find that the experimentally determined *τ*_*cat*_ across strains is best modelled by a gamma distribution 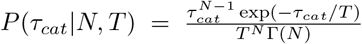, characterized by two parameters - a step parameter (*N*) and time-scale parameter (*T*), where Γ (*N*) is the usual Gamma function (Fig. 7b) (Gardner et al. 2011, Odde et al. 1995). We utilize the framework of Bayesian inference to deduce the parameters of MT-catastrophe distribution in different strains (see section A.1). Bayes rule assigns the posterior probability of a set of model parameters (***θ*** = {*N, T*}) given the observations (**O** = *τ*_*cat*_) by (Gelman et al. 2013),

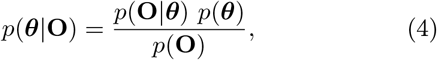

where *p*(***θ***) is the prior distribution of the model parameters, *p*(**O**|***θ***) is the likelihood function of the observations **O** given the parameters ***θ*** and *p*(**O**) = ∫ *p*(**O**|***θ***)*p*(***θ***) *d**θ*** is the marginal likelihood distribution of the observations (see section A.1). To estimate an informative prior, we obtain a likelihood distribution from the rather abundant and reliable data on the Mal3 strain (Fig. 7b, also *Methods*) and combine it with a flat (uniform) prior (see section A.1 for details). We then use this informative prior to obtain the distribution of parameters that best describes the observations *τ*_*cat*_ in each strain. As a result, we find that with the exception of *mcp11* Δ, *mal3* Δ and *tip1* Δmutant strains (all regulators of MT growth dynamics), all other strains studied have very similar parameters associated with the *τ*_*cat*_ distribution (Fig. 7d). Additionally, a truncated distribution obtained by restricting the full distribution at experimentally observed minimum and maximum values of *τ*_*cat*_ suffices to explain the observed empirical distribution.

**Figure 7:**
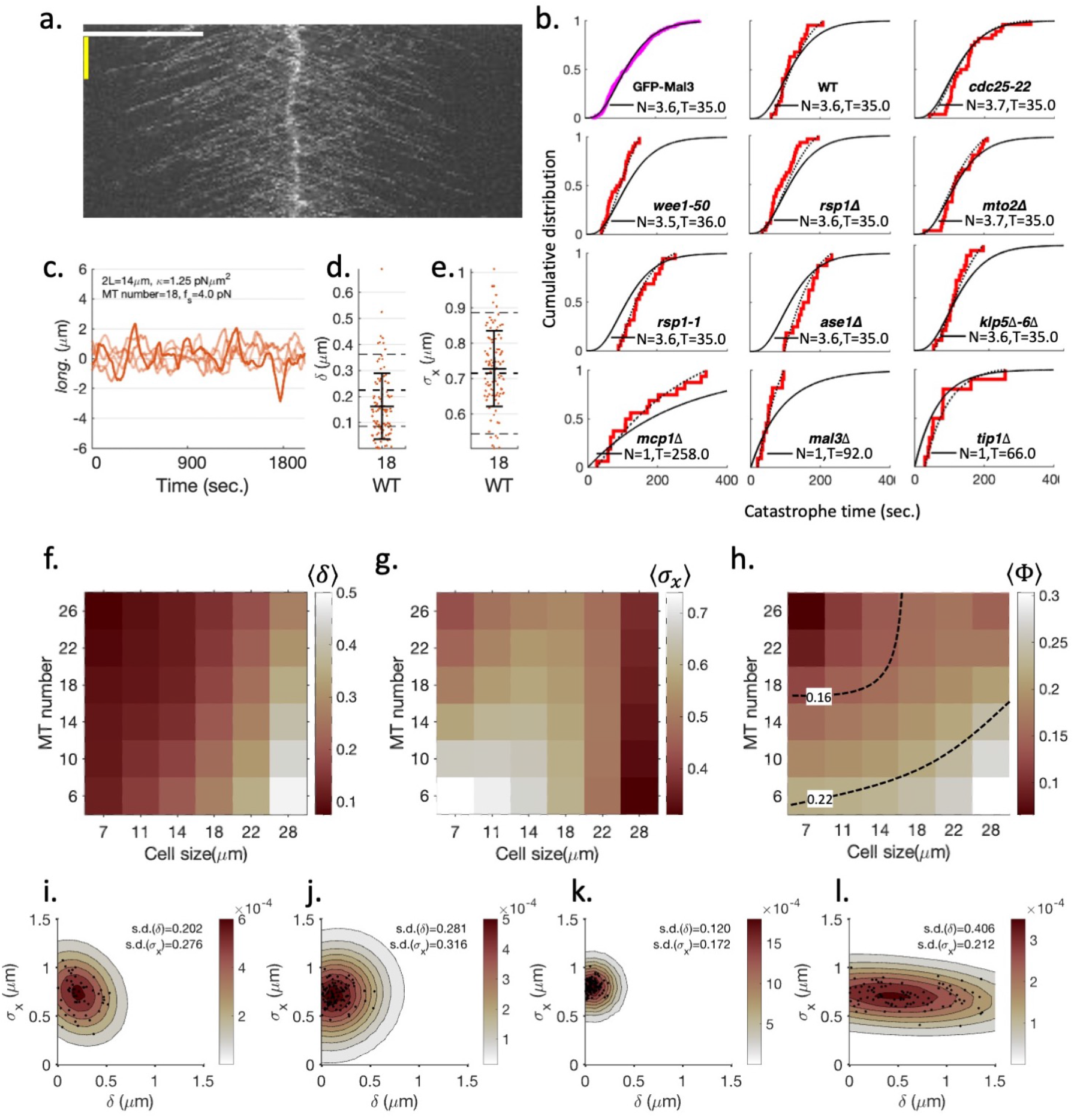
Stochastic model recapitulates optimality of WT. **a**. A representative kymograph of cell-wide projection of Mal3 strain (*cdc25-22* GFP-Mal3 Sid4-mCherry). We utilized the traces of comets with clearly visible ends to determine the catastrophe times (see *Methods*). Scale bars are 10 *µm* (white) and 120 s (yellow). **b**. (Top-left panel) Cumulative distribution of *τ*_*cat*_ (magenta) obtained from Mal3 strain. A gamma distribution gives the best fit (black curve) for the distribution of *τ*_*cat*_. (Other panels) Parameters obtained using the Bayesian analysis of the *τ*_*cat*_ obtained for various strain utilizing EnvyGFP-atb2 Sid4-mCherry background. The red line shows the cumulative distribution from experiment. The black curve shows the full distribution inferred using Bayesian analysis. The dotted curve is truncated distribution obtained using the estimated parameters (black curve) with truncation at the minimum and maximum of experimentally observed *τ*_*cat*_ for respective strains. **c**. Example trajectories of SPB obtained from simulations. **d.-e**. Statistics of *δ* (d.) and *σ*_*x*_ (e.) obtained using the parameters listed in (c.). The dotted lines are experimental values of means (wide-lines) and standard deviation (thin-lines) for respective statistics. Error bars are standard deviation. **f.-g**. Discrete contour-plots showing the systematic effect of variation in cell-length and number of MTs on *δ* (f.) and *σ*_*x*_ (g.). Color-bar scale is in *µm*. The rest of the parameter-set are same as in (c.). **h**. A discrete contour-plot showing the effect of variation in cell-length and number of MTs on failure-coefficient of SPB localization Φ. The dashed curves are iso-contours for two values of Φ. **i.-l**. Bivariate probability distribution of *δ* and *σ*_*x*_. Scatter plots show cell-to-cell variations. The contour show probability distribution estimated by fitting a 2D-Gaussian after convolving the data points with a symmetric Gaussian kernel. Color bar scale shows probability density. **i**. Experimental data from WT strain showing cell-to-cell variation in *σ*_*x*_ is larger than *δ* suggesting that the control on *δ* is *stiffer* than on *σ*_*x*_. **j**. Simulations with parameters listed in (c.) with the cell length following the experimentally observed cell length variation. **k**. Simulations with perfect correlation between left (*l*) and right (*r*) MT force generators, in terms of their local orientations, and iMTOC localization on the nucleus. **l**. Simulations with uncorrelated left-right MTs orientation.

We now proceed to estimate the remaining MT parameters, the stall force *f*_*s*_ and the flexural rigidity *k* (the growth and shrinkage velocities have been measured directly from experiments). We do this by fitting the simulated distributions of *δ* and *σ*_*x*_ from the model Eqs. 1, 2 (see SI) to the distribution obtained from the WT data (see Fig. S10). This gives a value of *k* ≈ 1.25p*Nµm*^2^, which goes to set *f*_*e*_. This value of *k* falls in the lower end of the range of MT stiffness obtained in *in vitro* measurements (Kikumoto et al. 2006), and is consistent with the bent and curved MTs observed in the WT as well as the small cell strains, such as the *wee1-50*. The fit however does not fix *f*_*s*_ as along as *f*_*s*_ > 1*pN*. If we choose *f*_*s*_ ranging from 4 pN to 6 pN, then we find that the mean catastrophe times, dwell-times of MTs growth dynamics and auto-correlation function of nucleus also matches with the experimental observations. For this range of values of *f*_*s*_, we are indeed in the regime *f*_*s*_ > *f*_*e*_, where MT buckles before the MT growth stalls.

Having established the kinetic parameters of our model that take very similar values in the different strains (Table S2 in SI), we are now in a position to compare the nuclear trajectories and the statics of nuclear positioning, *δ* and *σ*_*x*_, from the model with those obtained from the experiments (Fig. 7c-e). The cell-strain dependent features that we vary in the model include the MT-number *N*^*r*/*l*^, the orientation pattern of the filaments and the cell length (Fig. 7f and g).

Our stochastic model recapitulates the observation that *δ* and *σ*_*x*_ show opposite trends upon changing MT number and orientation and cell length away from the WT. For instance, from our model we find that independent of MT-number, *δ* increases with an increase in the cell length away from WT. This is consistent with our experimental observations. Moreover, *δ* decreases as a function of increasing MT-number, keeping everything else fixed.

On the other hand, the model shows that *σ*_*x*_ decreases with an increase in cell-size away from WT and decreases with increasing MT-number, again consistent with experimental observations. For small MT-number, the relative fluctuations of the force bearing MTs to the left and right is large, leading to an increase in *σ*_*x*_ (Dogterom & Yurke 1998, Howard 2006). Note that the MT-number affects both the MT-polymerization induced forces and the effective drag on the nucleus.

Our stochastic model recapitulates the observation that it is the combination of *δ* and *σ*_*x*_ that is functionally relevant for the position of the nucleus at the onset of mitosis. As before, this can be expressed by the mis-centered fraction of SPB localization Φ (Fig. 7h), with the following results: (i) Keeping cell length fixed (say, = 14 *µ*m), the value of Φ decreases monotonically as the MT-number increases before saturating at high MT-number. This suggests that a further increase in MT numbers does not translate into significant decrease in Φ, consistent with experimental findings (Sect. 2.3) (ii) Keeping MT-number fixed (say, = 18), the value of Φ increases beyond a certain cell length = 14 *µ*m (Fig. 7h). (iii) Along a fixed contour of Φ (say, Φ = 0.16 dashed curve in Fig. 7h), variation of the cell length away from WT (= 14 *µ*m) is compensated by larger MT numbers.

The anisotropy in the joint distribution of *δ* and *σ*_*x*_ provides on the *control* of nucleus positioning. The bivariate distribution width is much broader in the *σ*_*x*_ direction than the *δ* direction (Fig. 7i), implying that control on *δ* is *stiffer* than *σ*_*x*_ (Gutenkunst et al. 2007, Daniels et al. 2008). We find that the reliability *δ* is predominantly controlled by the strong correlation between the left and right oriented MT force generators and correlations in the localization of iMTOCs on the nucleus. On the other hand, the robustness *σ*_*x*_ is controlled by cell length, MT length scale and number of MT bundles. Figure 7j shows a comparison of the distribution of *δ* and *σ*_*x*_ between the experimental WT and the stochastic model where the cell lengths take values over the experimental range. In addition, in our model we can vary the left/right correlation of the MT force generators. We find that a perfect correlation reduces the width of the variation along *δ*, while a weaker correlation increases it substantially (Fig. 7k, l).

## 3 Discussion

In this paper, we have used live-cell imaging to study the fidelity of nucleus centering (and hence of the septum) in wild type (WT) and genetically perturbed fission yeast cells in terms of two phenotypic objectives for nucleus position at the onset of mitosis, namely reliability (*δ*), and robustness (*σ*_*x*_). The genetic perturbations that we have studied go to change cell length, MT-number and their orientational patterning and MT dynamics. In conjunction, we have analysed a stochastic model that describes the nucleus centering in terms of MT-polymerization induced forces. Extracting the parameters of this model from experiments, we were able to compute the corresponding *δ* and *σ*_*x*_, as one varied the MT-number, orientation and cell length. Both experiment and theory, show that high fidelity in septum centering is a consequence of the collective optimization of *δ* and *σ*_*x*_ in the nucleus positioning. Our analysis has interesting implications that we list below.

### 3.1 Optimal MT length and implications for cell length selection

The first implication is that the length scale of MTs growth needs to be comparable with the cell length. MT length-scale is determined by the distribution of catastrophe time along with the growth and shrinkage velocities. We showed that catastrophe time distribution across many perturbations can be modelled by a gamma distribution. The surprising finding is that the parameters of the distribution do not show significant difference, suggesting that MT dynamics and consequently the MT-length is unaltered by the perturbations we have imposed on the cells. This is likely because the perturbations do not affect the MT (dis)assembly machinery; specific MT regulator deletion strains do produce deviations in the MT-catastrophe times distribution. While it is possible that this distribution gets altered because of force-enhanced catastrophe (Janson et al. 2003), this appears to be small. This optimality in nucleus centering in the WT, as a consequence of the matching of the MT-length (in the milieu of the yeast cell) and the WT cell length, might have implications for cell-size selection in fission yeast.

### 3.2 Proper MT number and orientation for optimal force patterning

The proper centering of the nucleus is contingent on proper nonequilibrium force patterning that is generated by MT-polymerizing forces. Thus the force patterning is described by the orientation of the MTs, their arrangement in bundles and the number of MT-bundles. In fission yeast this is achieved by a noncentrosomal MT-organization with MT arranged in a small number of bundles emanating from the nucleus. Our study shows that the optimal orientation of MTs is when they form bundles coaligned with the long axis of the cell. Deviation from coalignment leads to aberrant centering. Any change from this arrangement increases *δ* and *σ*_*x*_. Organizing MTs in coaligned 3-4 bundles distributed around the nucleus diminishes transverse fluctuations *σ*_*y*_ driven by the torque.

The performance of the two phenotypic objectives improves with an increase in coaligned MTs (Dogterom & Yurke 1998, Howard 2006, Howard & Garzon-Coral 2017), however beyond a certain number, there is no significant gain. Beyond approximately *N* = 22 MTs, the improvement if any is marginal, suggesting as in any biological control system, a tradeoff between effectiveness and economy.

### 3.3 Combined optimization of *δ* and *σ*_*x*_ implies feedback and adaptation of microtubule pushing forces

The optimality of nucleus position fluctuations, codified by *δ* and *σ*_*x*_, is a natural consequence of the fluctuating MT-polymerization forces. The opposing MT-polymerization based pushing forces can be viewed as a stochastic control on a marginally stable nucleus confined within the cell.

We note that both in the experiments and the stochastic model of the WT, the cell-to-cell variation in robustness (*σ*_*x*_) is much larger than in reliability (*δ*), suggesting that *δ* is under stiff control by the cell, while *σ*_*x*_ is under a sloppier control (Gutenkunst et al. 2007, Daniels et al. 2008, Machta et al. 2013). This, while expected, deserves further attention. The control of *σ*_*x*_ is effected by the force-dependent catastrophe of MTs that we have included in our stochastic model - this acts as an adaptive feedback control, and possibly operates over a certain range of cell lengths. On the other hand, cellular control of *δ* is achieved by ensuring left (*l*) - right (*r*) symmetry in the MT-polymerizing forces. At present we do not have a complete understanding of the molecular basis of this force correlation, which will likely involve anti-parallel crosslinkers such as Ase1 (Janson et al. 2007) and motor proteins such as Kinesin-14 or Dynein (Janson et al. 2007, Carazo-Salas & Nurse 2005, Tan et al. 2018), an investigation of this cellular control mechanism is a task for the future.

## Methods

### Genetics, cell culture, and strains

Standard yeast genetics methods are followed to insert fluorescent markers and to create strains (Hagan et al. 2016). The EnvyGFP plasmid was a gift from Bi Lab (University of Pennsylvania). We use Gibson assembly to construct a pFa6a based plasmid to endogenously tag Atb2 with EnvyGFP inserted at the N-terminal and transformed it to the fission yeast cell. The list of primers used in this construct is given in Table S3. A list of the strains used in the work can be found in Table S4. Cells were maintained at 25°C on agar plates containing YE5S media. For microscopy-based experiments, a small number of cells were inoculated in liquid YE5S medium a night before the experiment and incubated at 25°C under shaking. The cells were harvested when they reached an optical density of ∼ 0.5 + 0.2 a.u. for microscopy experiments.

### Microscopy and image analysis

Imaging was performed using the spinning disk confocal microscope. Briefly, a Nikon Eclipse Ti2 inverted microscope, equipped with a Nikon CFI Plan Apochromat 100*x*/1.4 NA objective lens, a Nikon Perfect Focus System (PFS), a Mad City Labs integrated Nano-View XYZ micro- and nano-positioner, a Yokogawa Spinning Disk CSU-X1 unit, a Photometrics Evolve EM-CCD camera, controlled by Molecular Devices MetaMorph 8.0. For GFP and mCherry imaging, solid-state lasers of 488 nm (100mW) and 561 nm (100mW) were used. We utilized the CherryTemp microfluidics-based thermostat to precisely control the temperature while imaging to minimize the experimental variations and analyze 30-50 cells in each case. The samples were mounted on YE5S in 2% agar immobilized directly on a ready-made microfluidics chip-based set-up (CherryTemp, CherryBiotech) to control the temperature of the cell during microscopy (Velve Casquillas et al. 2011). Once we set up the temperature using the cherry temp on the microscope, we waited for ∼ 30 mins for cells to acclimate under the new condition before proceeding to image them. The exception was the microscopy of temperature-sensitive mutants *wee1-50* and *cdc25-22*, when we waited for ∼ 2 hours before imaging. The microscopy was performed at 35°C. The specific conditions for each kind of microscopy experiment are given below. All perturbations are imaged and analyze identically.

### Live cell microscopy and segmentation of SPB

A z-stack (total 7 focal planes spaced 1*µm* apart) were acquired in brightfield(Exposure time 20 ms), GFP(Exposure time 100 ms, EM Gain 400), and mCherry(Exposure time 100 ms, EM Gain 400) channel. The stack was acquired for ∼ 1 hour with a 15s time interval.

The segmentation and analysis were done using semi-automated scripts written in ImageJ macros and MATLAB (Schindelin et al. 2012, MATLAB 2016). We use the stack of brightfield images to segment cells based on the shift in the intensity of the cell boundary (Lugagne et al. 2018). We track the cell by identifying the cell at time *t* + 1 based on maximum projection. This method for segmenting and tracking the cells was very robust, nonetheless, we checked the segmented cells and made manual corrections when an error was found. To segment the Sid4 signal, we first construct an image with maximum projection in the mCherry channel of the segmented cell images, remove background via thresholding, and find a 3×3 pixel window with the highest integrated intensity.

### Live cell microscopy and segmentation of Nucleus

Similar procedure as above, with the exception that the stack was acquired for ∼ 1 hour with 30s time interval.

The cell and SPB are segmented using the procedure given above. To segment the Cut11 signal, we first construct an image with maximum projection in the GFP channel of the segmented cell images, binarize images via selecting the pixel having top 5% intensities and selecting the largest object.

### Live cell microscopy and segmentation of MT in EnvyGFP-Atb2 strains

A z-stack (total 13 focal planes spaced 0.5*µm* apart) were acquired in GFP (Exposure time 100 ms, EM Gain 400), and mCherry (Exposure time 100 ms, EM Gain 400) channel. The stack was acquired for ∼ 10 mins with a 6 s time interval. We also captured the BF image of the first and last time-point. The cell is segmented via setting a threshold in the GFP channel images after background subtraction and taking a maximum projection. Following this, we constructed a color combined image stack using GFP and mCherry channels. The MT which emanate from the SPB are manually segmented using ImageJ (segmented line tool) (see Fig. S7) and analyzed using MATLAB scripts.

The number of MT bundles, total MT mass, and orientation are measured using static z-stacks (similar setting as above). The number of MT-bundles is counted manually. The quantification of total MT mass and orientation was done using the following steps. First, we segmented the MT cytoskeleton using the “Trainable Weka Segmentation” plugin in ImageJ (Arganda-Carreras et al. 2017). A manually curated data set was used to train the classifier to distinguish between MT-cytoskeleton and cytosol. The segmented probability histogram was bimodal and we use the value corresponding to the minima to segment the MT cytoskeleton and create a segmented mask. MT-mass is estimated by directly integrating the total intensity in a sum projection under the segmented mask. The orientations are calculated using the OrientationJ plugin in imageJ (Püspöki et al. 2016) under the segmented mask.

### Live cell microscopy and segmentation of MT in Mal3 strains

A z-stack (total 13 focal planes spaced 0.5*µm* apart) were acquired in GFP (Exposure time 100ms, EM Gain 400), and mCherry (Exposure time 100ms, EM Gain 400) channel. The stack was acquired for ∼ 10 mins with a 6s time interval. We also took the BF image of the first and last time-point.

For each cell, we constructed 2 types of kymographs (see Fig. S8). 1. A cell-wide and used it to score the catastrophe event in long MTs and 2. A kymograph using 2×50 pixel window centered around SPB was used to score events which happened in a short time-scale. In both cases, we manually traced the trajectory of the MT growth and analyze them in MATLAB.

### Microscopy and segmentation of septum

We incubated the cells at 35°C for 3 hours and than image them for quantifying the septum positioning. As known, the *cdc25-22* cell does not initiate the mitotic phase at the restrictive 35°C. Thus, to quantify the septum position in *cdc25-22* cells, we incubate the cells at 35°C for 3 hours and than quench the temperature to 25°C. Most of the cell undergo mitosis immediately and form a septum after 40 − 50 mins. To segment septum position, we manually segment the cell end and septum using the point-tool in ImageJ.

## Acknowledgements

We thank members of Tran lab at Institut Curie and members of the Simons Centre, NCBS-TIFR for fruitful discussions. IJ thanks Sergio A. Rincon, Ana Loncar, Lara K. Kruger, Manuel Lera Ramirez, Federica Arbizzani, Anne Paoletti, Vincent Fraisier, Ludovic Leconte, Amit Kumar, Raj Hosein, and Jyotsana J. Parmar for their support and discussion. We thank Mukund Thattai, Matthieu Piel, Vaishnavi Ananthanarayanan, Alkesh Yadav and Krishnan Iyer for comments and insights on the manuscript. IJ and MR acknowledge support from the Department of Atomic Energy (India), under project no. RTI4006, and the Simons Foundation (Grant No. 287975). MR acknowledges a J.C. Bose Fellowship from DST-SERB (India). IJ acknowledges a Curie-NCBS-InStem postdoctoral fellowship. PTT acknowledges grant support from INCa, Fondation ARC, and La Ligue National Contre le Cancer - Ile de France. PTT is a member of the Labex CelTisPhyBio, part of IdEx PSL. The authors greatly acknowledge the Nikon Imaging Centre @ Institut Curie-CNRS, member of the French National Research Infrastructure France-BioImaging (ANR-10-INSB-04), and computational facilities at NCBS.

## Notes

### Competing Interest Statement

The authors have declared no competing interest.

