## Supplementary Information for "Optimal reliability, robustness and control of nucleus centering in fission yeast is contingent on nonequilibrium force patterning"

### A Supplementary information

#### A.1 Determining MT-catastrophe time distribution using Bayesian inference

Arriving at the correct statistics of the MT growth dynamics is crucial for a description of the MT-driven processes. However, estimating MT growth dynamics by observing MTs (EnvyGFP-Atb2) has limitations – we can only reliably follow the growth dynamics of the longest MT emanating from a MT-bundle. This limits the length (and time) window to observe the MT's catastrophe events, and consequently, the measured catastrophe length or time distributions only represents a truncated subset of the entire distribution. Moreover, the lengths (and time) window for observing the MTs depend on the MT number, orientation and cell length, which may introduce systematic biases in the analysis, or may result in sparse statistics. Instead of using conventional approaches, such as maximum likelihood estimate (MLE), an alternative method is to ask how well the data explains a set of model parameters. This is achieved by the Bayesian inference method (Gelman et al. 2013). Bayes rule assigns the posterior probability on a set of distribution parameters ($\boldsymbol{\theta}$) given the observations ($\mathbf{O}$) by:

$$p(\boldsymbol{\theta}|\mathbf{O}) = \frac{p(\mathbf{O}|\boldsymbol{\theta})\ p(\boldsymbol{\theta})}{p(\mathbf{O})}, \tag{5}$$

where $p(\boldsymbol{\theta})$ is the prior distribution of the model parameters, $p(\mathbf{O}|\boldsymbol{\theta})$ is the likelihood function of the observations $\mathbf{O}$ given the parameters $\boldsymbol{\theta}$ and $p(\mathbf{O}) = \int p(\mathbf{O}|\boldsymbol{\theta})p(\boldsymbol{\theta})\, d\boldsymbol{\theta}$ is the marginal likelihood distribution of the observations. Apart from the data set of observations $\mathbf{O}$, we also require the following to evaluate the above expression: 1. a probability distribution function (e.g. exponential, gamma, etc.) which is parametrized by the set $\boldsymbol{\theta}$ and, 2. a distribution of prior probabilities of $p(\boldsymbol{\theta})$.

We use a gamma distribution, defined by a step parameter ($N$) and a time-scale parameter $T$, to describes the MT catastrophe time ($\tau_{cat}$) (Odde et al. 1995, Gardner et al. 2011, Duellberg et al. 2016):

$$P(\tau_{cat}|N,T) = \frac{\tau_{cat}^{N-1}\exp(-\tau_{cat}/T)}{T^N\Gamma(N)},$$

where $\Gamma(N)$ is the gamma function. At $N = 1$, the gamma distribution reduces to an exponential distribution, as has been considered previously (Foethke et al. 2009, Dogterom & Leibler 1993, Glunčić et al. 2015). We find that the gamma distribution is much more general, and hence more suitable to represent MT-regulator-driven changes in the distribution of catastrophe time. To validate the choice of the distribution, we use the GFP:Mal3 Sid4:mCherry *cdc25-22* (Mal3 strain) strain to measure the $\tau_{cat}$. By incubating the strain at the restrictive temperature (35°C), we can measure the MT +end dynamics in very long cells (see Fig. S8). The MT seldom reaches the tip of these cell, thus observation of MT dynamics is free from the effect of force. In our hand, the growth rate of MT is very similar to the growth rate observed in the EnvyGFP:Atb2 strains (WT). The cumulative distribution of catastrophe time is plotted in Figure 7c yields the best fit by a gamma distribution with $N = 3.6$ and $T = 35.0\,$s using Maximum likelihood estimate, in agreement with the reasoning presented above.

To construct the prior probability distribution $p(N,T)$ on the model parameters (i.e., $N$ and $T$), for the second part of the calculation, we use the following information. We assume that the catastrophe time distribution is established robustly by the cell, as we observe that the MT growth dynamics parameters are very similar across the many perturbations (see Table S1). Thus we use a prior biased by the parameters of the catastrophe time distribution seen in the Mal3 strain. To accomplish this, we combined the likelihood distribution for catastrophe time data from the Mal3 strain and a uniform distribution (a flat prior). The prior takes the form $p(N,T) \sim W_1 * p(N,T|O) + W_2 * U$, where $p(N,T|O)$ is the likelihood distribution evaluated from Mal3 strain data ($O$) (see Fig. S8), $U$ is a uniform distribution and $W_1$ and $W_2$ are weights. The weights are selected to ensure that the ratio of maximum and minimum probabilities in $p(N,T)$ is $\approx 10$ (see Fig. S8). However, a small deviation from this ratio does not lead to different results. We evaluated the posterior distribution $p(N,T|\mathbf{O})$ using the grid method on the support $N \in [1:10], T \in [10s:500s]$. The evaluated $p(N,T|\mathbf{O})$ for all the strains are given in Fig. S9. Even with this bias in the prior distribution, the data from *mcp1*Δ, *mal3*Δ and, *tip1*Δ show the highest value of $p(N,T|\mathbf{O})$ at $N \sim 1$ (i.e. an exponential distribution), assuring that the method indeed allow deviations in the predicted parameters. In Fig. 7, we also show the full distribution made using the value of $N$ and $T$ at their respective $max(p(N,T|\mathbf{O}))$. Most interestingly, a truncated distribution made using the predicted parameters and with the experimentally observed support (i.e. $\tau_{cat} \in [a,b]$, where $a$ and $b$ respectively are minima and maxima of the observed $\tau_{cat}$) shows an excellent fit with the experimentally observed distribution.

Table S1: **MT-growth dynamics parameters.** p-values are calculated with respect to WT using Wilcoxon rank sum test. $***$ for $p \leq 10^{-4}$, $**$ for $p \leq 0.005$, $*$ for $p < 0.05$, ns is not significant.

| Strain | $V_o^+$ $(\mu m/min)$ | $V_o^-(\mu m/min)$ | $V_f^+$ $(\mu m/min)$ | $\langle\tau_{cat}\rangle\pm$ S.D. (sec.) | $\langle\tau_{dwell}\rangle\pm$ S.D. (sec.) | n |
|---|---|---|---|---|---|---|
| WT | $4.8 \pm 1.4$ | $-13.8 \pm 5.9$ | $3.0 \pm 0.9$ | $115.5 \pm 36.9$ | $45.5 \pm 36.0$ | 22 |
| *cdc25-22* | $4.4 \pm 1.3$(ns) | $-15.5 \pm 4.2$(ns) | $2.9 \pm 1.6$(ns) | $148.4 \pm 62.2$( $*$) | $17.3 \pm 25.0$( $**$) | 28 |
| *wee1-50* | $5.0 \pm 1.9$(ns) | $-11.2 \pm 4.4$(ns) | $2.9 \pm 1.6$(ns) | $90.2 \pm 33.1$( $*$) | $54.7 \pm 32.7$(ns) | 32 |
| *rsp1*Δ | $5.4 \pm 1.5$(ns) | $-13.3 \pm 4.9$(ns) | $3.1 \pm 1.0$(ns) | $95.7 \pm 38.5$(ns) | $40.8 \pm 30.4$(ns) | 29 |
| *mto2*Δ | $6.9 \pm 3.4$( $**$) | $-10.9 \pm 6.0$(ns) | $3.2 \pm 1.6$(ns) | $126.5 \pm 45.4$(ns) | $64.4 \pm 40.3$(ns) | 27 |
| *rsp1-1* | $4.4 \pm 2.4$(ns) | $-16.7 \pm 6.2$(ns) | $2.8 \pm 0.8$(ns) | $154.8 \pm 46.4$( $*$) | $63.2 \pm 39.4$(ns) | 19 |
| *ase1*Δ | $4.3 \pm 1.2$(ns) | $-13.2 \pm 5.8$(ns) | $2.5 \pm 0.9$(ns) | $157.9 \pm 41.1$( $**$) | $51.8 \pm 32.7$(ns) | 16 |
| *klp5*Δ-*klp6*Δ | $5.3 \pm 1.7$(ns) | $-12.7 \pm 4.4$(ns) | $3.0 \pm 1.3$(ns) | $107.8 \pm 31.8$(ns) | $45.1 \pm 33.4$(ns) | 21 |
| *mcp1*Δ | $5.3 \pm 3.2$(ns) | $-11.8 \pm 5.2$(ns) | $3.9 \pm 4.7$(ns) | $152.0 \pm 101.8$(ns) | $64.3 \pm 66.4$(ns) | 16 |
| *tip1*Δ | $4.2 \pm 1.6$(ns) | $-12.7 \pm 5.3$(ns) | NA | $88.2 \pm 71.4$( $*$) | NA | 10 |
| *mal3*Δ | $4.2 \pm 0.9$(ns) | $-11.6 \pm 7.3$(ns) | NA | $52.2 \pm 23.6$( $***$) | NA | 11 |

#### A.2 Stochastic model for MT-dependent nucleus centering

Here we discuss a model for microtubule (MT) pushing-force dependent nucleus centering to help us understand the diversity of phenomena observed in our experiments. The framework takes into account the motion of the nucleus because of stochastic forces arising due to MT polymerization against the cell wall. The model also accounts for the non-linear growth dynamics of MT, number, and orientation of MT, etc., inside the cell. Since the imaging plane is an x-y projection, we will take the cell to be a 2-dimensional rectangle of length $2L$ and width $2R$ (see Fig. S10a). The nucleus is modeled as a rigid circle, and MT nucleation sites are situated at MTOCs located at the periphery which nucleates MTs with swivel points.

As a consequence, the nucleus undergoes translational dynamics according to,

$$\frac{d\boldsymbol{X}}{dt} = \frac{1}{\zeta^T}\left\{\sum_{i=1}^{N_r}\boldsymbol{f}_i^r(\psi_i,\theta_i,t) + \sum_{i=1}^{N_l}\boldsymbol{f}_i^l(\psi_i,\theta_i,t) + \boldsymbol{f}_{th}\right\}, \tag{6}$$

where $\boldsymbol{X} = [\mathrm{x},\mathrm{y}]$ is the position vector of the centroid of the nucleus, and $\zeta^T$ is the hydrodynamic translational drag tensor (see below). On the right hand side, $\boldsymbol{f}_i^r(\psi_i,\theta,t)$ and $\boldsymbol{f}_i^l(\psi_i,\theta,t)$ are the forces due to the $i^{th}$ microtubule emanating toward the right $(r)$ or the left $(l)$ directions respectively, located at an angle $\psi$ with respect to the SPB $(\psi_{SPB} = 0)$ and with an orientation $\theta$ with respect to the long axis of the cell. $N_r$ and $N_l$ are the number of MT-nucleation sites on the nucleus aligned towards the right or left sides respectively at time $t$. $\boldsymbol{f}_{th}$ are random forces of thermal origin, with statistics $\langle\boldsymbol{f}_{th}\rangle = 0$ and $\langle\boldsymbol{f}_{th}(t)\cdot\boldsymbol{f}_{th}(t')\rangle = 6k_bT\zeta^T\delta(t-t')$.

The polymerizing MTs also induce rotational dynamics on the nucleus,

$$\frac{d\boldsymbol{\omega}}{dt} = \frac{1}{\zeta^R}\left\{\sum_{i=1}^{N_r}\boldsymbol{T}_i^r(\psi_i,\theta_i,t) + \sum_{i=1}^{N_l}\boldsymbol{T}_i^l(\psi_i,\theta_i,t) + \boldsymbol{T}_{th}\right\}, \tag{7}$$

where $\boldsymbol{\omega}$ is the angle between the longitudinal axis of the cell and vector the connecting center of the nucleus to SPB (denoted as $\boldsymbol{r}$). $\zeta^R$ is the hydrodynamic rotational drag tensor (see below). Akin to the forces, $\boldsymbol{T}_i = \boldsymbol{r}_i \times \boldsymbol{f}_i$ is the torque applied by $i$th MT on the nucleus where $\boldsymbol{r}_i = r_{nuc}\hat{\mathbf{n}}$, where $r_{nuc}$ is the radius of the nucleus and $\hat{\mathbf{n}}$ is the unit vector in the direction of $i^{th}$ MTOC. $\boldsymbol{T}_{th}$ are random torques of thermal origin, with statistics $\langle\boldsymbol{T}_{th}\rangle = 0$ and $\langle\boldsymbol{T}_{th}(t)\cdot\boldsymbol{T}_{th}(t')\rangle = 6k_bT\zeta^R\delta(t-t')$.

Before we proceed, we need to specify the form of the hydrodynamic drag tensors $\zeta^T, \zeta^R$ that enters into the dynamical equations above. We first argue that the dominant mechanism of momentum dissipation in fission yeast is via cytoplasmic viscosity and not, as in mammalian cells, the actin cortex viscosity (Rupprecht et al. 2018). This is because the actin cortex in fission yeast cell is very sparse (Mishra et al. 2014) and unlike the mammalian cell there is no connected actomyosin mesh during interphase. Furthermore, the smallest gap between the nuclear membrane and the cortex is around $0.3\,\mu$m. The drag arising from the cytoplasmic viscosity is an effective drag that has contributions from both the nucleus and attached MTs. The cytoplasmic drag is likely enhanced due to the presence of intracellular structures such as the ER. Despite the complex nature of the cytoplasmic fluid, no spatial dependence has been observed in the dynamics of small lipid droplets in fission yeast cytoplasm (Selhuber-Unkel et al. 2009), suggesting that the yeast cytoplasm behaves like an isotropic Stokesian fluid. Consistent with this, experiments on tagged particle dynamics show the usual diffusive behaviour $R^2 \sim t^\alpha$, with $\alpha$ distributed close to 1 (with a median = 0.8) (Selhuber-Unkel et al. 2009, Tolić-Nørrelykke et al. 2004). We note that recent experiments (Molines et al. 2022) suggest that MT dynamics is strongly influenced by cytoplasmic viscosity, reinforcing our claim that the effective drag must have contributions from both the nucleus and the MT.

Since the dissipation from the nucleus and the MTs (taken as dashpots) appear in series, their individual translational drag coefficients combine as,

$$\frac{1}{\zeta^T} = \frac{1}{\zeta^T_{nuc}} + \frac{1}{\zeta^T_{MT}} \tag{8}$$

while the rotational drag predominantly has contributions from the rotating nucleus alone. The drag coefficients of the nucleus (considered as a rigid sphere of radius $r$) in the cytoplasm of viscosity $\eta$ is isotropic,

$$\zeta^T_{nuc} = 6\pi\eta r \tag{9}$$

and

$$\zeta^R_{nuc} = 8\pi\eta r^3 \tag{10}$$

The translational drag coefficients of the MTs is anisotropic, with its parallel and perpendicular components given by

$$\zeta^{T,\parallel}_{MT} = \sum_i \frac{2\pi\eta l_i(t)\cos(\theta_i)}{\ln(l_i(t)\cos(\theta_i)/2r_{MT}) - 0.2} \tag{11}$$

and

$$\zeta^{T,\perp}_{MT} = \sum_i \frac{4\pi\eta l_i(t)\sin(\theta_i)}{\ln(l_i(t)\sin(\theta_i)/2r_{MT}) + 0.84} \tag{12}$$

where $l_i(t)$ is the length of the longest MT emanating from the nucleus perimeter from angle $\psi_i$ and $r_{MT}$ radius of the MT (Howard & Hyman 2003). The expressions Eqs. 11 and 12 are valid in the infinite rod limit, i.e. when $\frac{r_{MT}}{\langle l \rangle} \ll 1$.

The above forms of the hydrodynamic drag are approximations to the real system. Both the finite geometry of the yeast cell that confines the cytoplasmic fluid and the permeability of the nucleus as it moves through the incompressible cytoplasmic fluid, affect the form of the drag. Denoting the permeability of the nucleus by $k$, the drag coefficient reduces by $\zeta_{nuc} = \frac{6\pi\eta r_{nuc}}{1+k/2r^2_{nuc}}$ (Joseph & Tao 1964). On the other hand, the drag on the nucleus increases because of geometric confinement. Treating the yeast cell as a cylinder of radius $R$, with the assumption of axial symmetry, the drag coeffcient follows $\zeta^T_{nuc} = \frac{9\pi^2\sqrt{2}\eta r_{nuc}}{4(\frac{R-r_{nuc}}{R})^{5/2}}$, evaluated to the lowest order in $\frac{R-r_{nuc}}{R}$ (Happel & Brenner 1983). Finally, we have treated the nucleus as a rigid sphere. The deformability of the nucleus can also change the coefficient of drag.

We have experimentally estimated the translational diffusion of the nucleus (using MBC treated cells) and found the translational diffusion length scale to be of order $10^{-2}\mu m$ which is significantly smaller than the variation of the nucleus position because of active forces (which are of order $10^0\mu m$). Consequently, we neglect the effect of thermal forces in Eq. 6 and Eq. 7.

The microtubules apply forces only when they are in contact with the cell boundary and this is given by

$$\boldsymbol{f}^r_i(\psi_i, \theta_i, \mathrm{t}) = \begin{cases} \boldsymbol{f}_p & \text{if } f_p \le f_e \\ \boldsymbol{f}_e, & \text{otherwise} \end{cases} \tag{13}$$

where $\boldsymbol{f}_p$ is the pushing force applied by MTs because of polymerization when MTs are in contact with the cell cortex and $\boldsymbol{f}_e$ is the critical buckling force. $\boldsymbol{f}_p$ is given by:

$$\boldsymbol{f}^l_p = -\mathrm{f_s}(1 - \frac{\mathrm{V}^+_f}{\mathrm{V}^+_o})\hat{\mathbf{n}}\Theta(\mathrm{x(t)} + r\cos(\omega + \psi) - \mathrm{l}_i(\mathrm{t})\cos(\omega_i + \psi_i + \theta_i) - L) \tag{14}$$

assuming that the applied force prominently affects the 'on' rate (the probability of a tubulin subunit to get intercalate at the tip of microtubule) of tubulin subunits (see Peskin 1993, Dogterom 1997). Here $\mathrm{f_s}$ is the stall force (i.e. force at which the MT growth halts), $\mathrm{V}^+_o$ and $\mathrm{V}^+_f$ are the mean growth rate of MTs during the free-growth phase and during the contact-phase with cell boundary (while MTs applying pushing force), $\mathrm{l}_i(t)$ is the length of the MT. $\hat{\mathbf{n}} = [\cos(\theta_i), \sin(\theta_i)]$ is the unit orientation vector of the MT. The Heaviside theta function $\Theta(\mathrm{x(t)} + r\cos(\omega + \psi) -$

Table S2: **List of model parameters and their reference values**

| Parameter | Definition | Reference value |
|---|---|---|
| $2L$ | Length of the cell | Variable (for WT=$14\mu m$ see Fig. 2) |
| $2R$ | Width of the cell | $3.2\mu m$ |
| $\psi_i$ | Angle of $i$th MTOC | Random uniform distribution |
| $\theta$ | Orientation of MT | Random with statistics of MT orientation in WT cells |
| $r_{nuc}$ | Radius of the nucleus | $1.3\mu m$ |
| $\eta$ | Viscosity of the cytoplasm | $\approx 0.9pNs/\mu m^2$ (Molines et al. 2022, Tolić-Nørrelykke et al. 2004, Foethke et al. 2009) |
| $r_{MT}$ | Radius of the MT | $0.013\mu m^a$ |
| $N = N_r + N_l$ | Number of MTs | 18 (for WT cells (Höög et al. 2007)) |
| $f_s$ | Stall force for MT growth | Fitted |
| $\kappa$ | Effective flexural rigidity of MTs | Fitted |
| $V_o^+$ | Growth velocity of MTs | Table S1 |
| $V_o^-$ | Shrinkage velocity of MTs | Table S1 |
| $V_f^+$ | Growth velocity of MTs during dwell phase | Table S1 |
| $P(\tau_{cat})$ | Distribution of catastrophe times | See Fig. 7 |
| $\Delta T$ | Time-step | 1 sec. |

[a] In our calculations, each bundle is made of upto 6 MTs, which may have a large radius toward the core of the bundle. However, small variation in the $r_{MT}$ doesn't affect the results.

$l_i(t)\cos(\omega_i + \psi_i + \theta_i) - L)$ enforces the condition that microtubules only apply force when they are in contact with the cell boundary. $\boldsymbol{f}_e$ is the critical buckling force require to buckle MT with flexural rigidity $\kappa$ and is given by:

$$\boldsymbol{f}_e = max \begin{cases} \kappa\pi^2/l_i(t)^2 \\ \kappa\pi^2/l_{io}^2 \end{cases} \tag{15}$$

where $l_{io}$ is the length of MT which can be constrained inside the geometry of the cell after buckling. The inclusion of $l_{io}$ emulate constraints imposed by cell geometry on the configurations of buckled MTs.

The growth dynamics of microtubule is stochastic where microtubules show dynamic instability. During the growth phase, MT polymerize with rate $V_o^+$ and during the shrinking phase, MT depolymerize with rate $V^-$. The microtubule will catastrophe (i.e. jump from the growing to shrinking phase) such that the catastrophe time $\tau_{cat}$ has a distribution given by $P(\tau_{cat})$. We also model force dependent reduction of catastrophe time. Here, if there is no force applied on MT then the catastrophe time $\tau_{cat}$ has a distribution given by $P(\tau_{cat})$. In the presence of force, the catastrophe time is given by $\tau_{cat}(V_+^f) = \tau_o + \frac{\tau_{cat}(V_+)-\tau_o}{V_+}V_+^f$, where $\tau_o$ is the mean catastrophe time at $V_+ = 0$ (Janson et al. 2003, Foethke et al. 2009). The value of $\tau_o$ is not known *in vivo*, but it is found to be $\sim 25$ sec in the reconstitution experiments (see (Janson et al. 2003)).

We use the Monte-Carlo integration approach to get a time-resolved solution to the above model. The approach has 2-steps: 1. Monte Carlo step– which evolves the growth dynamics of MT. 2. Numerical integration step – which updates the position of the nucleus. In the Monte Carlo step, either MTs length changes depending on their growth state or MTs switch from growing to shrinking state or vice versa. The switch from growing to shrinking state follow the distribution $P_e = 1 - \exp(-\Delta t/\tau_T)$, where $\tau_T = \frac{1-\int_0^{\tau_{cat}=T} P(t')dt'}{P(\tau_{cat}=T)}$ is the mean catastrophe time of MT with age $T$ (Gardner et al. 2011), and $\Delta t$ is the time-step. The switch from shrinking state to growing state occurs instantaneously if the $l_i <= 0$. In the Numerical integration step, we update the position of the nucleus using the forward Euler method for explicit integration. We initiate the calculation by assigning a fixed number of MT-anchoring sites which are distributed on the nucleus randomly. The number of nucleation sites (i.e. number of bundles) is equal to $((N_r + N_l) \div 4)$ (i.e. quotient of the fraction). The MTs are then randomly assigned to these sites and the MT-anchoring site with the maximum number of MT is considered the SPB to defining $\psi_{SPB}$. MTs orientation $\theta$ follow the experimentally determined orientation statistics of MTs and each time MTs switch from the shrinkage to growing state a new $\theta$ is assigned to an MT. The $\theta$ of the longest MT originating from the each anchoring site is considered in evaluation of the drag forces. Throughout this study, we assumed $N_r = N_l$ (unless stated otherwise). We started each simulation with the nucleus located on the tip of the cell and performed a simulation for $\approx 7200$ s (a generation time of fission yeast cell), unless noted otherwise.

We utilize $P(\tau_{cat})$ obtained from Mal3 strain (Cdc25-22 GFP-Mal3 Sid4-mCherry) and anticipated that the difference in catastrophe times seen in experiments can be attributed entirely to the force dependent change in velocities. Thus the model has only two free parameters - $\kappa$ and $f_s$. We systematically vary these parameters to search for a set of conditions which matches the experimental observations (see Fig. S10). We see that that for $\kappa \approx 1.25 pN\mu m^2$ the value of $\sigma_x$ matches with experimental observations, attributed to the dominance of buckling and $f_s$ only becomes a limiting parameters. For $f_s$, ranging from $4-6$ pN, the $\langle \tau_{cat} \rangle$ and dwell-times also matches with the experimental observations.

Table S3: **List of DNA primers used for constructing EnvyGFP:Atb2:KanMX6 plasmid**

| Name | Sequence |
|---|---|
| pfa6a-fwd | ATTCATCGATGATATCAGATCCACtagt |
| pfa6a-rev | TTAATTAACCCGGGGATCCGTC |
| patb2-fwd | gatccccgggttaattaaAGCATTTCCAGTCATATTTAACTC |
| patb2-rev | ctttagacatTTCGAACGTCTTTCCAGGGT |
| envy-fwd | gttcgaAATGTCTAAAGGCGAGGAATTGTT |
| envy-rev | atggatccTTTGTACAATTCGTCCATTC |
| atb2-fwd | attgtacaaaGGATCCATGAGAGAGATCATTTCCATTCACGTTGGC |
| atb2-rev | tccatgtcGCTGGCCGGGTGACCCGG |
| ter-fwd | cggccagcGACATGGAGGCCCAGAATAC |
| ter-rev | ctgatatcatcgatgaatAACCGACAACTCAGTAGTCTTATT |

Table S4: **List of strains**

| | |
|---|---|
| TP5070 | h- Cut11:GFP:Ura Sid4mC:Nat |
| TP4070 | h- ENVY-atb2:KanMx |
| TP4623 | h+ Sid4mC:Nat |
| TP4628 | h- Sid4mC:Nat Eatb2:Kan |
| TP4878 | h- Eatb2:Hph |
| TP4880 | h- Eatb2:Hph Sid4mC:Nat |
| TP4756 | h- Eabt2:Kan Sid4mC:Nat Cdc25-22 |
| TP4855 | h- Eatb2:Kan Sid4mC:Nat Wee1-50 |
| TP4763 | h- Eabt2:Kan Sid4mC:Nat Rsp1D:Hph |
| TP4889 | h+ Eatb2:Hph Sid4mC:Nat Mto2D:kan |
| TP4758 | h- Eabt2:Kan Sid4mC:Nat Rsp1-1 |
| TP4629 | h+ Sid4mC:Nat Eatb2:Kan Ase1D:Hph |
| TP4640 | h? Sid4mC:Nat Eatb2:Kan Klp5D:Ura Klp6D:His |
| TP5056 | h- Eatb2:Hph Sid4mC:Nat Mcp1D:Kan |
| TP4635 | h+ Sid4mC:Nat Eatb2:Kan Mal3D:Hph |
| TP5058 | h- Eatb2:Hph Sid4mC:Nat Tip1D:Kan |
| TP5066 | h? KanMx-pnmt1-GFP-Mal3 Sid4mC:Nat Cdc25-22 |

#### B Supplementary figures

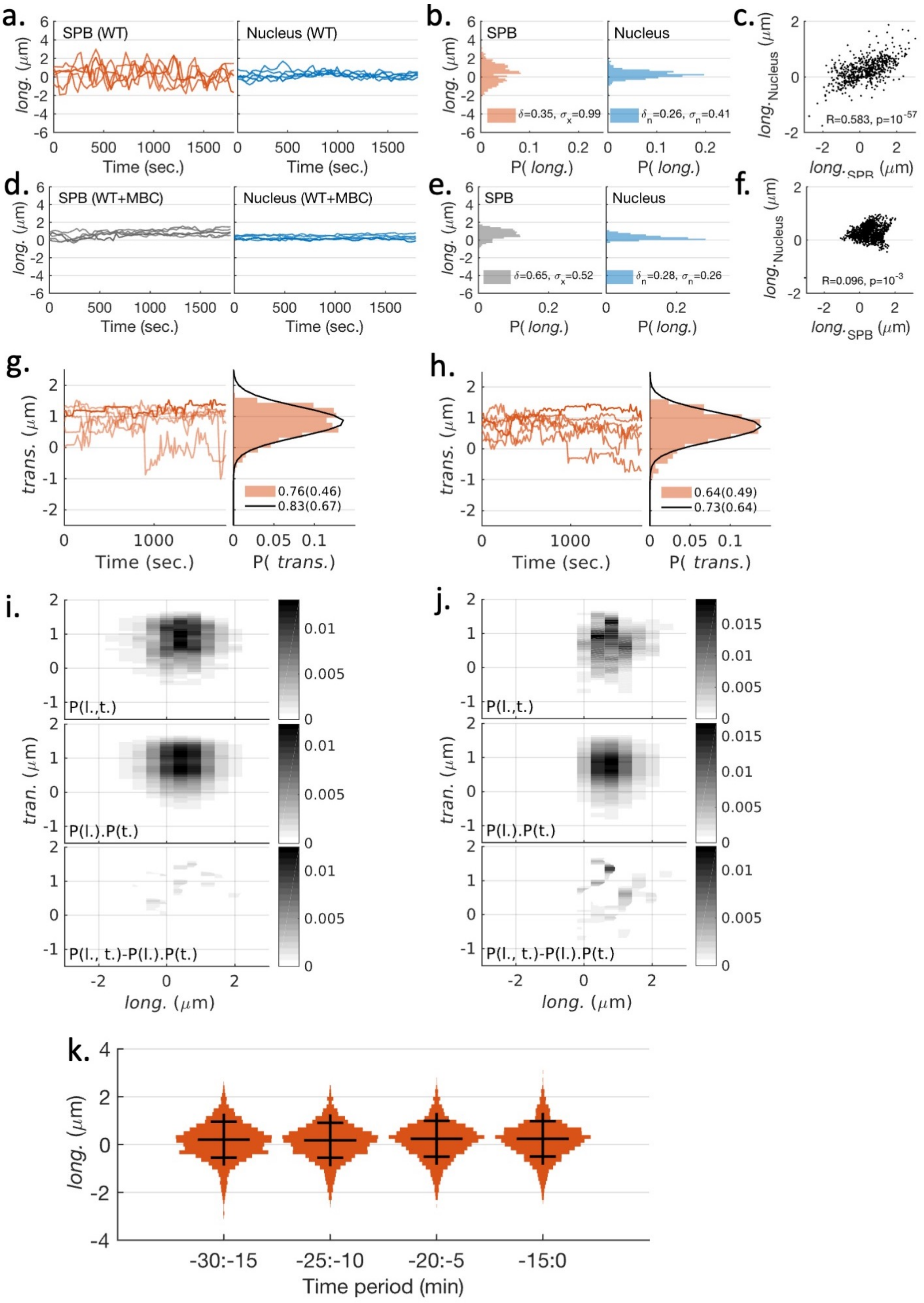


Figure S1: Nucleus centering in WT cells. **a.-d.** Few examples of longitudinal displacement of SPB (left) and centroid of the nucleus (right) w.r.t. the center of the cell in WT and MBC treated (WT+MBC) cells. Nucleus fluctuations vanish upon MBC treatment (as seen previously, (Tran et al. 2001)). **b.-e.** The distribution of longitudinal positions of SPB (left) and nucleus centroid (right) in WT (e) and MBC treated cells (h). **c.-f.** Covariance between the nucleus and SPB position for WT (c) and WT+MBC condition (f). **g.-h.** Examples of transverse displacement of SPB (left) and their distribution (right) in WT-cell (b) and with MBC treatment (e), respectively. **i.-j.** Joint distribution of the longitudinal and transverse position of SPB (top), the multiplication of marginal distribution of longitudinal and transverse position calculated independently (middle), and the subtraction of the two (bottom) for WT (c) and MBC treated (f) cells. These suggest that longitudinal and transverse positions are statistically independent, i.e., knowledge about the longitudinal position does not provide information about the transverse position and *vice versa*. **k.** Time-series of SPB dynamics before the onset of mitosis is stationary. Violin plot of the distribution of SPB-positions at different time-periods (x-axis label) before mitosis. Error bars are standard deviation.

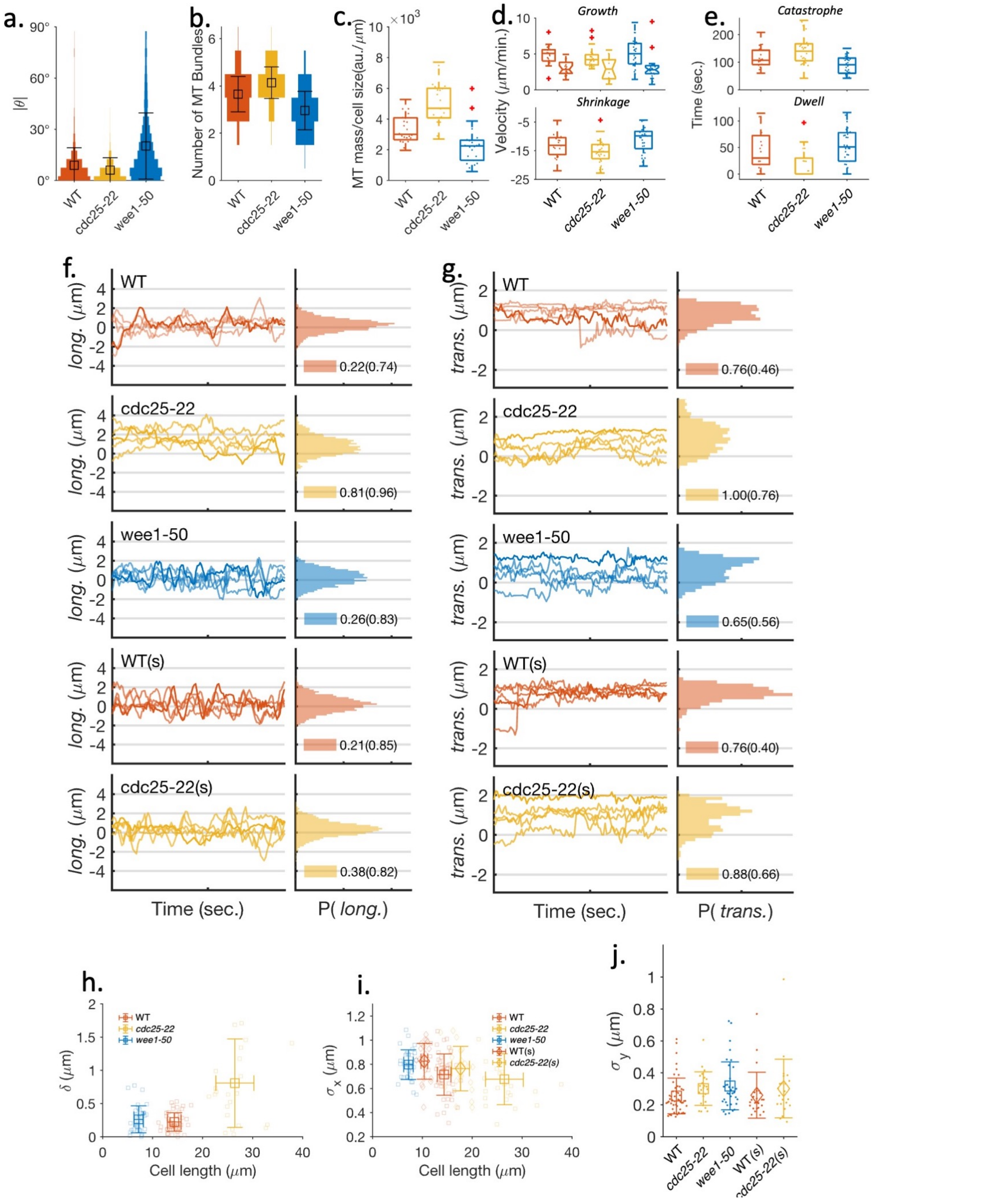


Figure S2: **a.-c.** Characterization of MT organization (also Fig. S6). **a.** The distribution of the local MT orientation $\theta$, the local angle of MT filament relative to the longitudinal axis of the cell (error bars are SD). **b.** Distribution of the number of MT bundles with means equal to 3.7 (WT), 4.1 (*cdc25-22*), and 3 (*wee1-50*) (error bars are SD). **c.** MT-mass, measured from the intensity of MT fluorescence, scaled with cell length. **d.-e.** MT growth-dynamics parameters. **d.** Distributions of free growth velocity (top, boxed), growth velocities on contact (top, notched), and shrinkage velocity (bottom), are very similar across the cell strains (depicted by corresponding colors; see Table S1 in SI). **e.** The distribution of catastrophe times (top) and dwell times (bottom): show a dependence on cell length. **f.** Examples of time series of longitudinal SPB displacements relative to the cell center in different strains and conditions (left) and the histograms of longitudinal SPB positions in the population (right) (legend show means and (s.d.)). **g.** The representative trajectories of transverse fluctuations and histogram of transverse SPB positions (legend show means (s.d.)). Transverse fluctuation usually remains bound within a narrow range. Exceptionally we also see large, sudden movement. **h.-i.** Scaling properties of $\delta$ (f) and $\sigma_x$ (g) with the cell length. **j.** Standard deviation in transverse displacement ($\sigma_y$) of SPB estimated for each cell. $\sigma_y$ does not show any correlation with cell length.

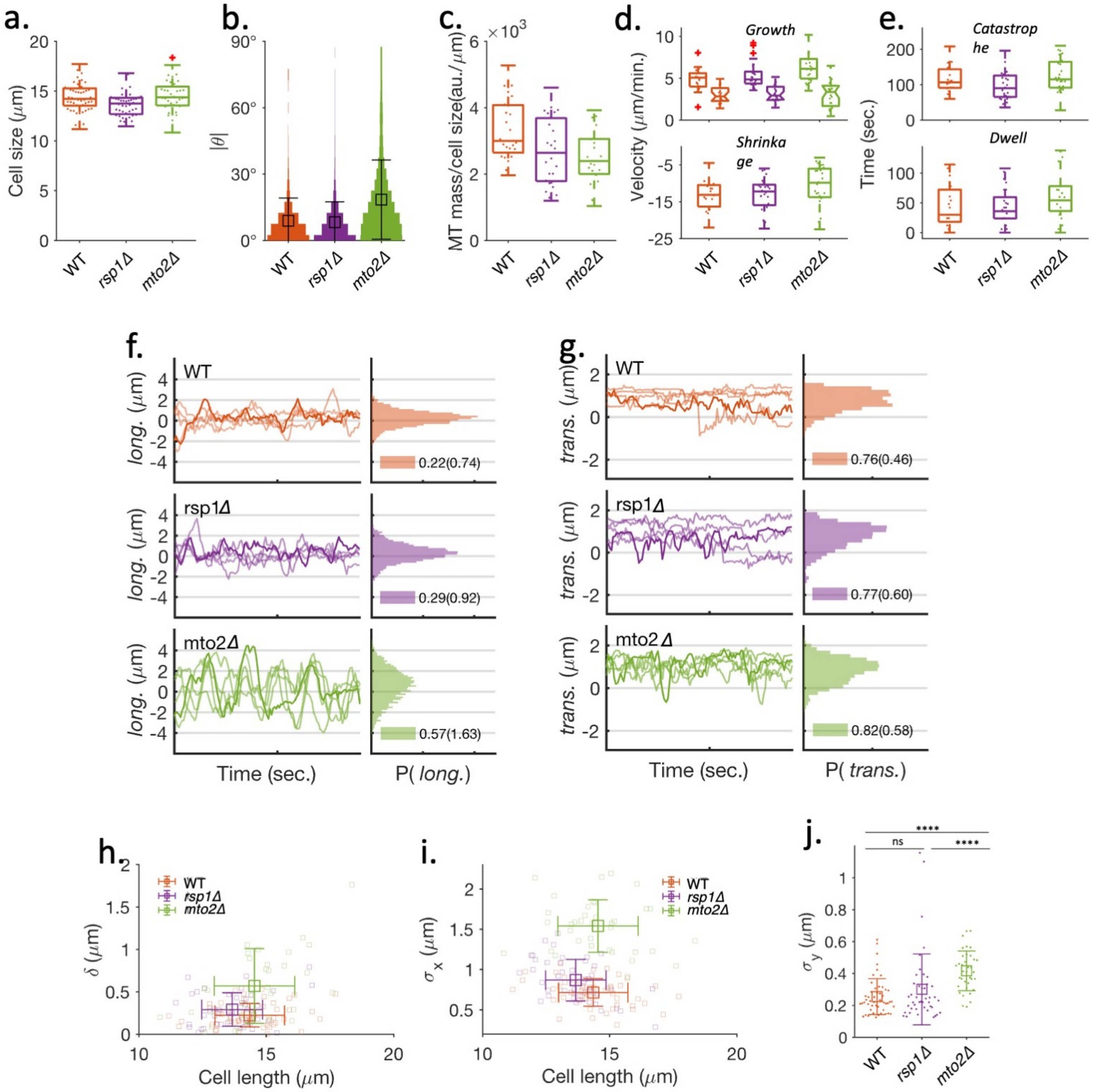


Figure S3: **a.** Cell length distribution at the onset of mitosis (also Fig. S6). **b.** The distribution of the local MT orientation $\theta$, the local angle of MT filament relative to the longitudinal axis of the cell (error bars are SD). **c.** MT-mass scaled with cell length, measured from the intensity of MT fluorescence. **d.-e.** MT growth-dynamics parameters. **d.** Distributions of MT growth velocity (top, boxed), MT growth velocities with contact (top, notched), and shrinkage velocity (bottom), for the different strains, are depicted by corresponding colors. (see Table S1 in SI). **e.** Distribution of catastrophe times (top) and dwell times (bottom). **f.** Examples of time series of longitudinal SPB displacements relative to the cell center in different strains (left) and the histograms of longitudinal SPB positions in the cell population (right) (legend show means and (s.d.)). **g.** The representative trajectories of transverse fluctuations and histogram of transverse SPB positions (legend show means (s.d.)). **h.-i.** $\delta$ (h) and $\sigma_x$(i) plotted against the cell length for WT, *rsp1*$\Delta$, and *mto2*$\Delta$ strains. Changes in $\delta$ and $\sigma_x$ with respect to the variation in MT-bundle numbers are independent of cell length. **j.** Standard deviation in transverse displacement ($\sigma_y$) of SPB estimated for each cell. $\sigma_y$ is significantly large in *mto2*$\Delta$ cells.

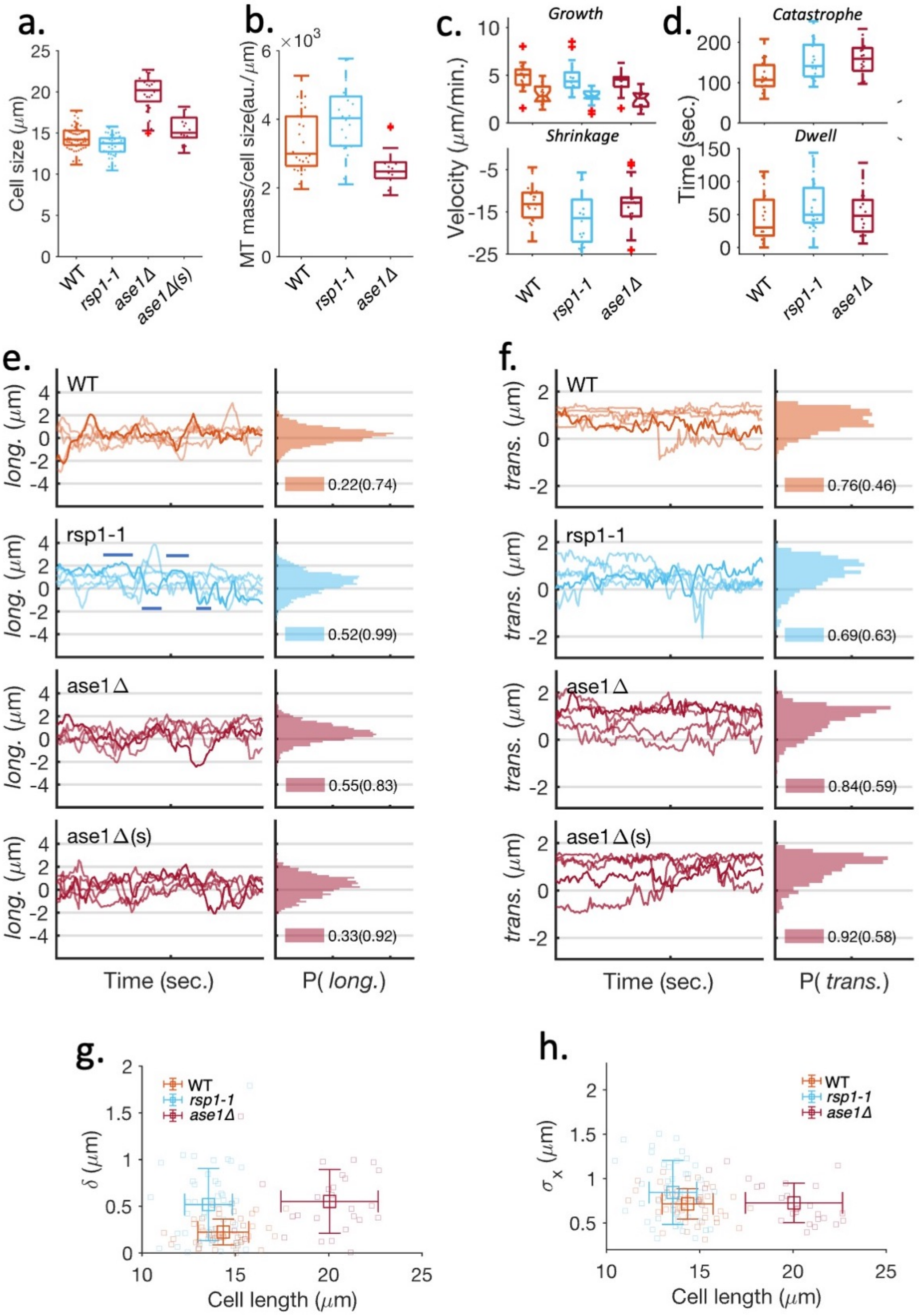


Figure S4: **a.** Cell length distribution at the onset of mitosis (also Fig. S6). **b.** MT-mass, measured from the intensity of MT fluorescence scaled with cell length. **c.-d.** MT growth-dynamics parameters. **c.** Distribution of MT growth velocity (top, boxed), MT growth velocities with contact (top, notched), and shrinkage velocity (bottom), depicted by corresponding colors (see Table S1 in SI). **d.** Distribution of MT catastrophe time (top) and dwell time (bottom). **e.** Examples of time series of longitudinal SPB displacements relative to the cell center in different strains and conditions (left) and the histograms of longitudinal SPB positions in the cell population (right) (legend show means and (s.d.)). In *rsp1-1* mutants, many cells show pauses in SPB dynamics when the SPB is poleward localized (highlighted by horizontal lines). These states mostly reflect highly asymmetric, aster-like MT arrangement (see Fig. 4**a.**, middle panel). **f.** The representative trajectories of transverse fluctuations and histogram of transverse SPB positions (legend show means (s.d.)). **g.-h.** $\delta$ (g) and $\sigma_x$(h) plotted against the cell length for WT, *rsp1-1*, and *ase1*$\Delta$ strains.

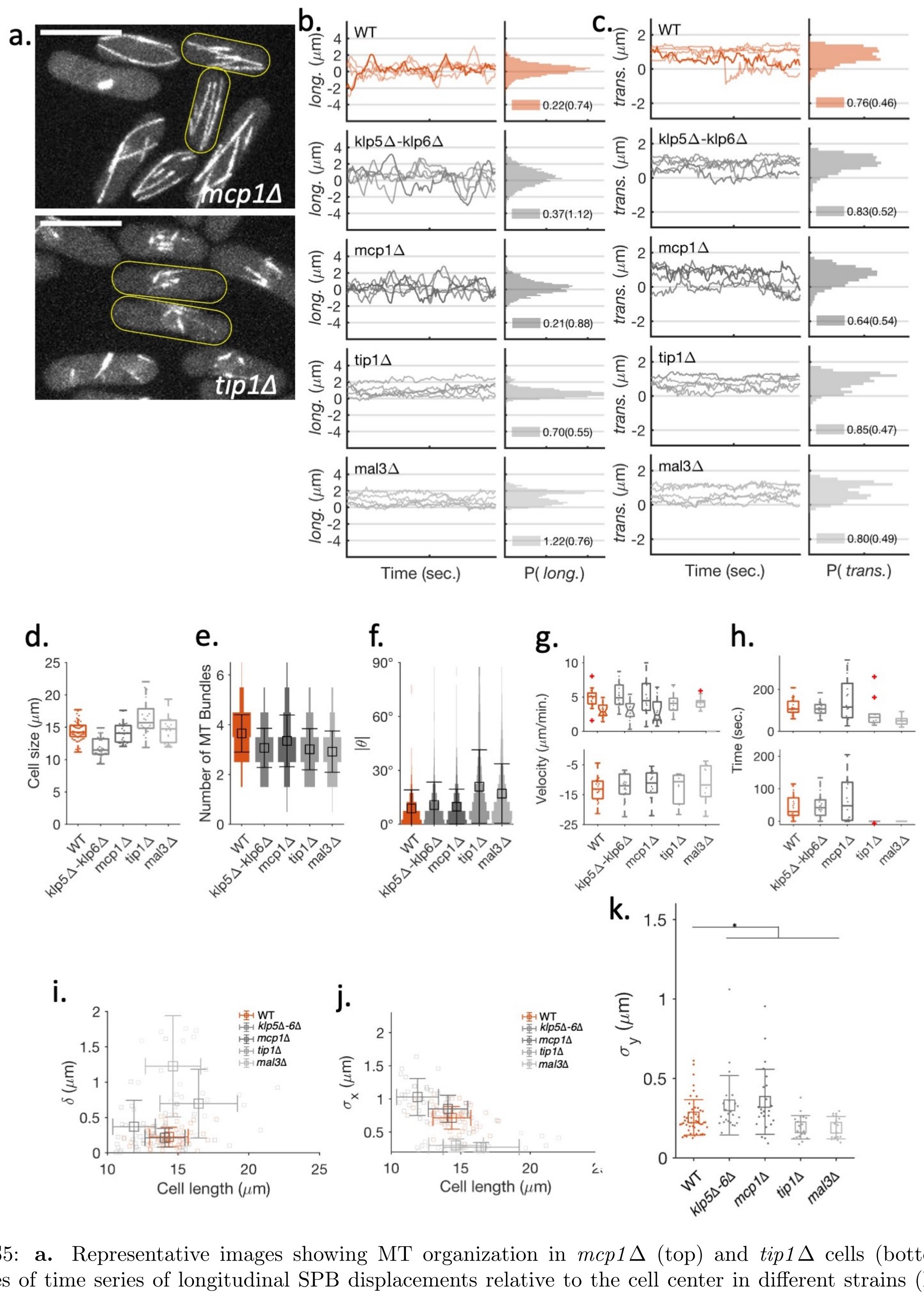


Figure S5: **a.** Representative images showing MT organization in *mcp1*Δ (top) and *tip1*Δ cells (bottom). **b.** Examples of time series of longitudinal SPB displacements relative to the cell center in different strains (left) and the histograms of longitudinal SPB positions in the cell population (right) (legend show means and (s.d.)). **c.** The representative trajectories of transverse fluctuations and histogram of transverse SPB positions (legend show means (s.d.)). **d.** Cell length distribution at the onset of mitosis (also Fig. S6). **e.** Distribution of the number of MT bundles (error bars are SD). **f.** The distribution of the local MT orientation $\theta$, the local angle of MT filament relative to the longitudinal axis of the cell (error bars are SD). **g.-h.** MT growth-dynamics parameters. **g.** Distribution of MT growth velocity (top, boxed), MT growth velocities with contact (top, notched), and shrinkage velocity (bottom), in the different strains, depicted by corresponding colors. **h.** Distribution of MT catastrophe time (top) and dwell time (bottom). **i.-j.** $\delta$ (h) and $\sigma_x$(i) plotted against the cell length for WT, *klp5*Δ-*klp6*Δ, *mcp1*Δ, *tip1*Δ, and *mal3*Δ strains. **k.** Standard deviation in transverse displacement ($\sigma_y$) of SPB estimated for each cell. $\sigma_y$ is significantly large in *klp5*Δ-*klp6*Δ and *mcp1*Δ strains and significantly small in *tip1*Δ and *mal3*Δ strains.

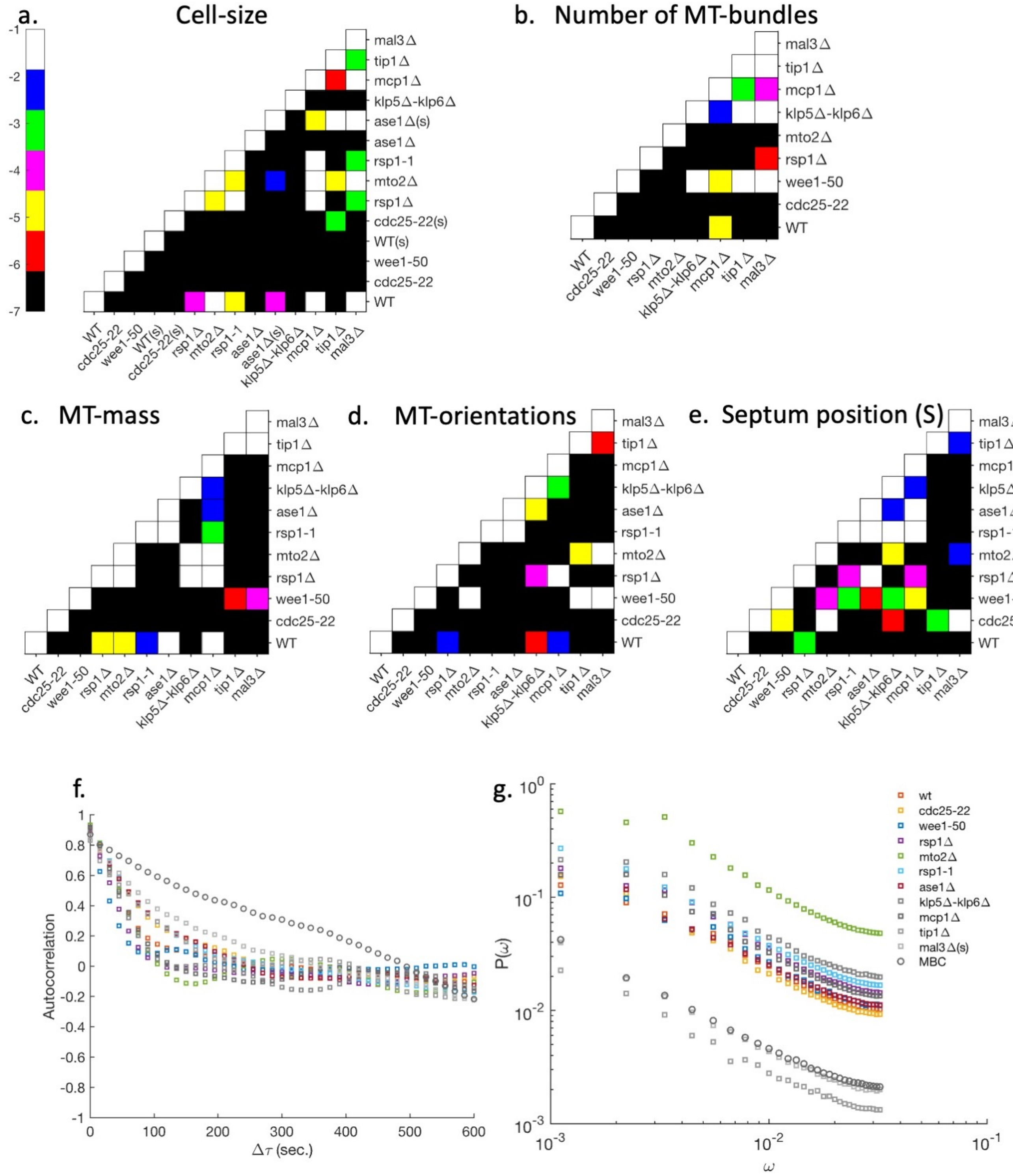


Figure S6: P-values were calculated using the rank-sum test across strains: **a.** cell length, **b.** the number of MT bundles, **c.** MT-mass, and **d.** MT-orientation. **e.** Septum position (S). The colobar shows the scale of p-values as the power of 10. **e.** Auto-correlation of longitudinal SPB position. **f.** Power spectrum of longitudinal SPB position. The power spectrum does not peak at any particular frequency, suggesting that the SPB dynamics does not have a defined period.

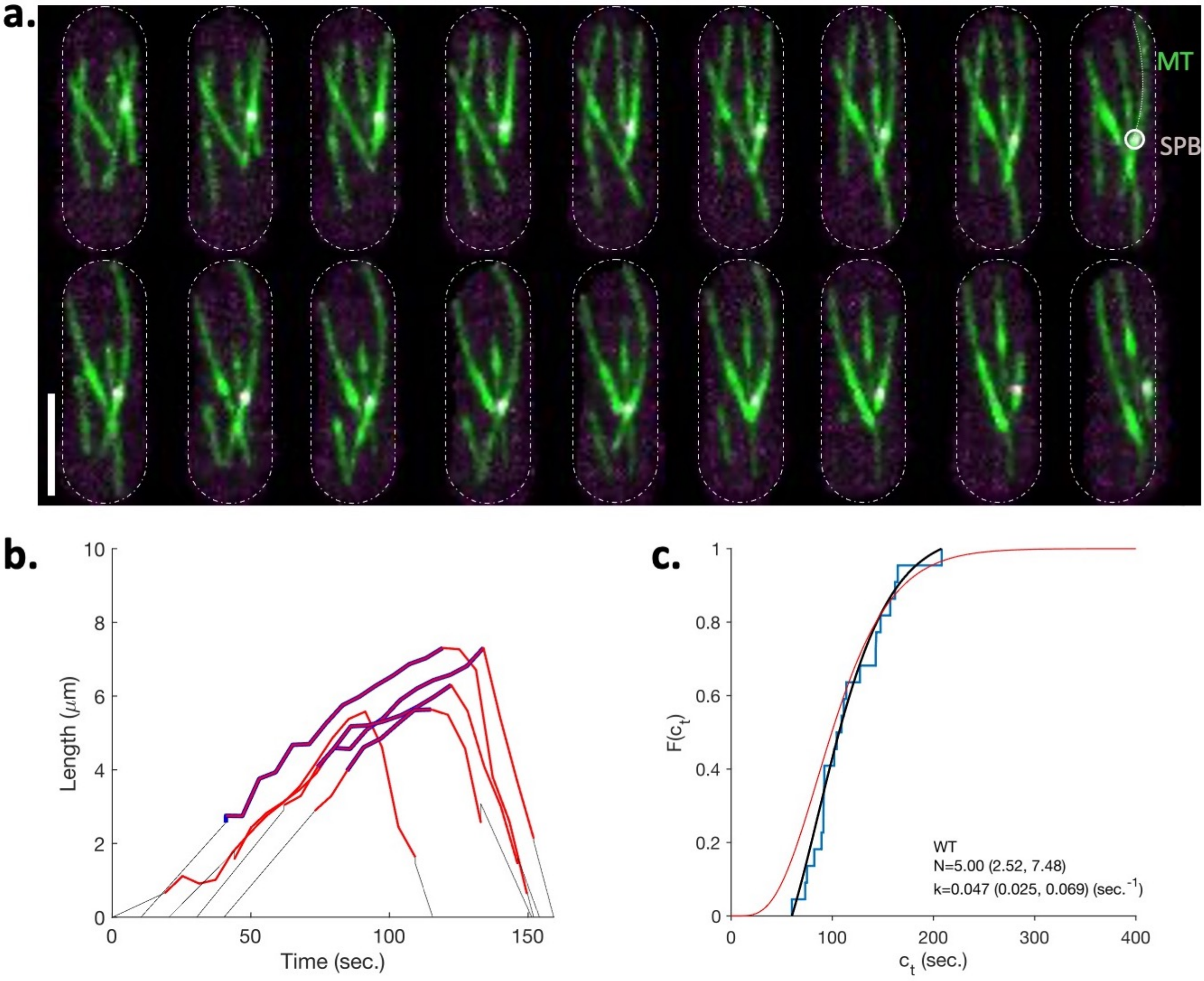


Figure S7: Measurement of MT growth dynamics parameters. **a.** A representative time-lapsed image sequence of a cell endogenously expressing EnvyGFP:Atb2 (MT, green) and Sid4:mCherry (SPB, cyan) in the WT background. Images are 6 seconds apart. While growing and making contact with the cell wall, MT show buckled morphologies and leads to large displacements in SPB. We traced the MT bundles length-wise to segment the growing MTs from a bundle originating from SPB. An example of such a trace is shown. **b.** Example of MT growth dynamics. Red traces show the measured length of MT from the SPB. The blue trace overlay represents the dwell phase. The dotted lines are an extrapolation of the growth trajectory evaluated using at least five first (for growth) and last (for shrinkage) data points for each MT. **c.** Empirical distribution function of catastrophe time observed in WT cells (blue curve). The black curve is fit using a maximum likelihood estimator using a truncated gamma distribution. The red curve shows the untruncated gamma distribution obtained using the fit parameters.

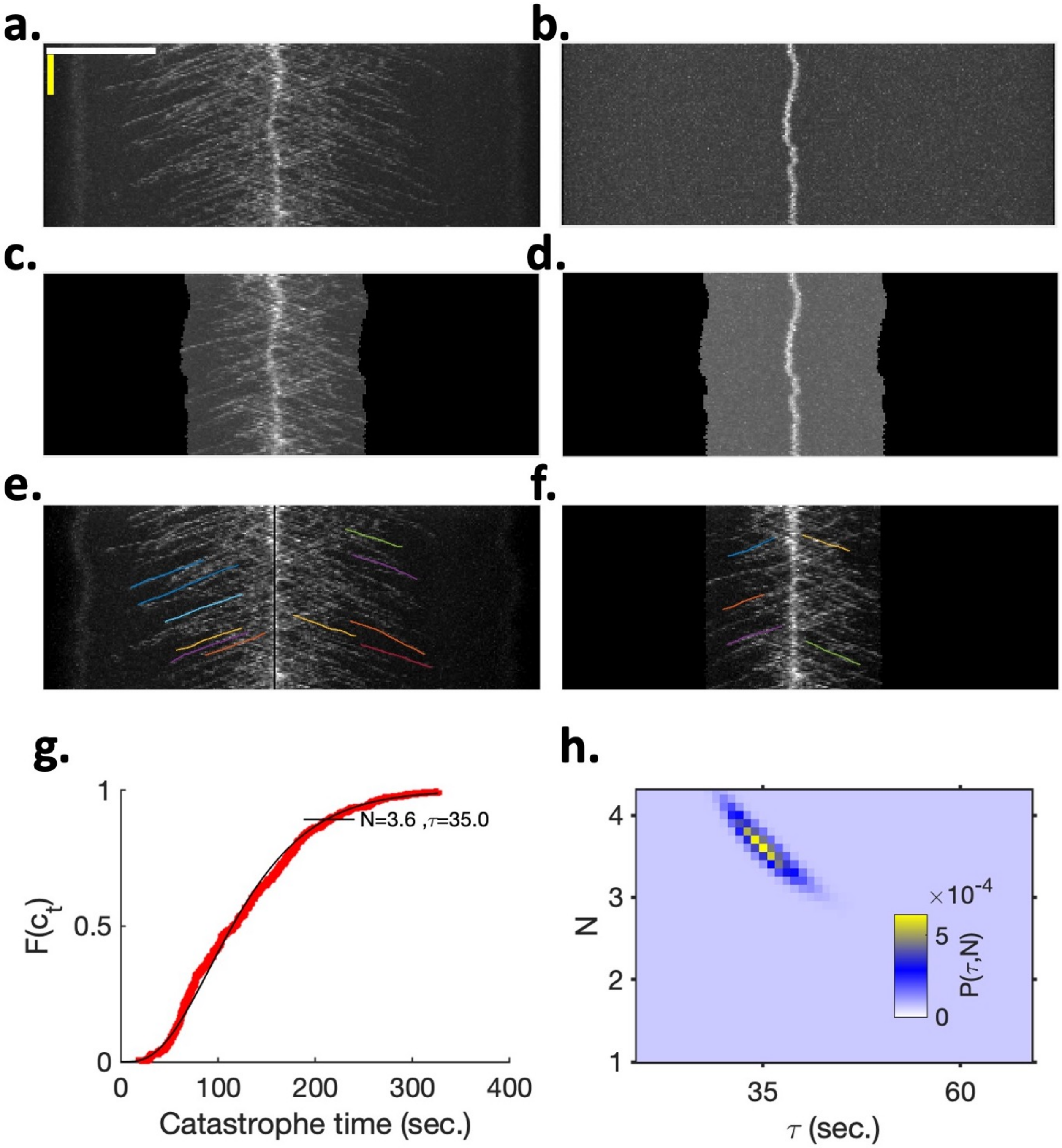


Figure S8: Catastrophe time distribution in long *cdc25-22* cells. **a.** Example of a whole-cell wide kymograph of Mal3 traces in a cell expressing GFP:Mal3 and Sid4-mCherry in *cdc25-22* background. The white scale bar is 10 $\mu m$ and the yellow scale bar is 120 sec. **b.** Sid4 (SPB) trace in the same cell. **c.-d.** Mal3(c) and Sid4(d) traces using a 2- by 50-pixel size rectangle around SPB. **e.-f.** Registered kymograph of Mal3 traces using SPB trace as reference. We manually segment only those Mal3 traces where the end is clearly visible. **g.** Red curve: Empirical distribution function of catastrophe times measure in long *cdc25-22* cells. The black curve shows the fitted gamma distribution using MLE. **h.** Informative prior used in Bayesian analysis of catastrophe times in various strains. The prior is constructed by combining a flat prior and the likelihood distribution derived from the Mal3 strain data set.

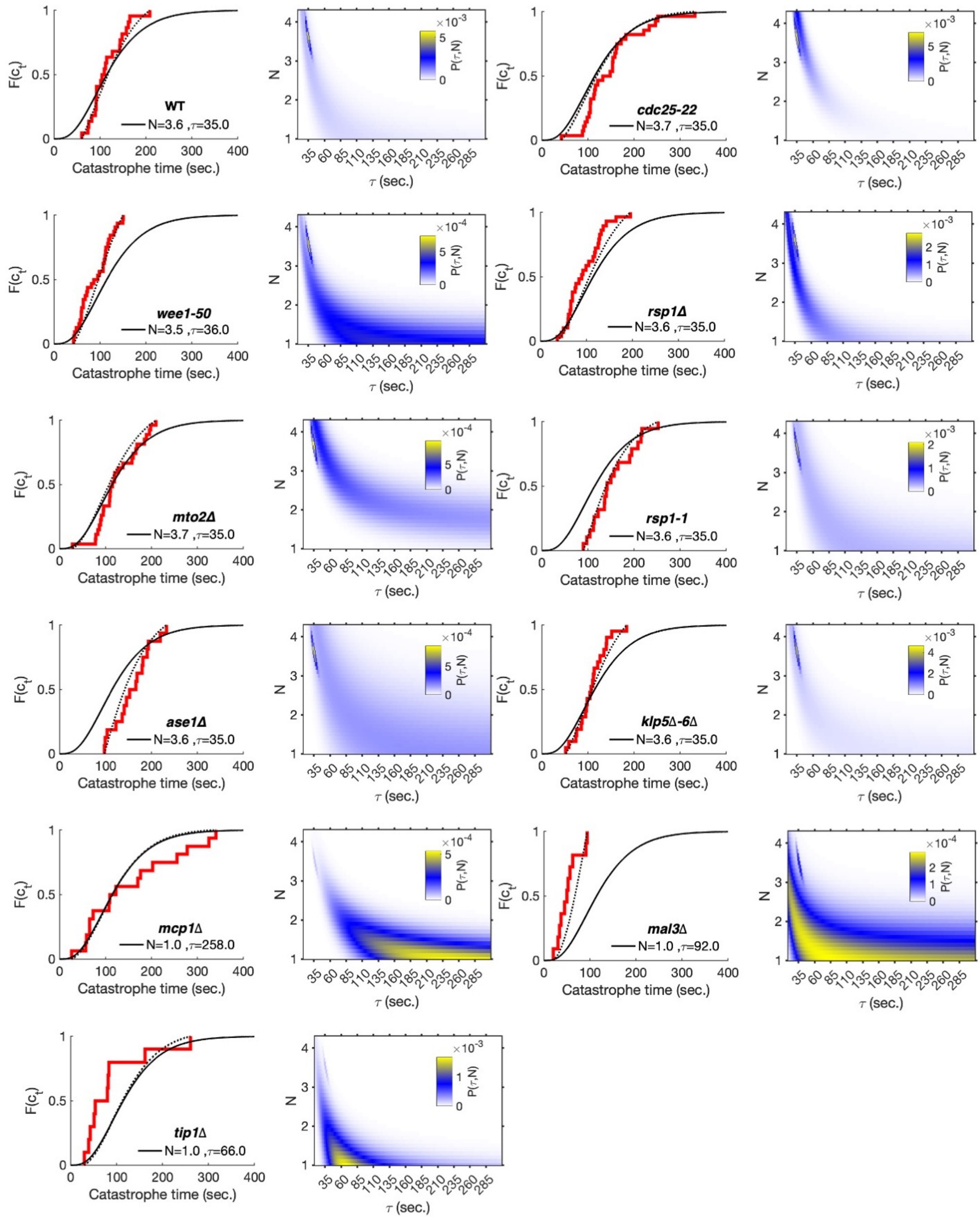


Figure S9: For each denoted strain: (left) Parameters obtained using the Bayesian analysis of the catastrophe times. The red line shows the empirical probability distribution. The black curve shows the full distribution. The dotted curve is truncated distribution obtained using the estimated parameters (black curve) with truncation at the minimum and maximum of experimentally observed catastrophe times for respected strains. (right) Joint probability distribution of step parameter (N) and time-scale parameter ($T$) using catastrophe time data from the observation MT growth dynamics.

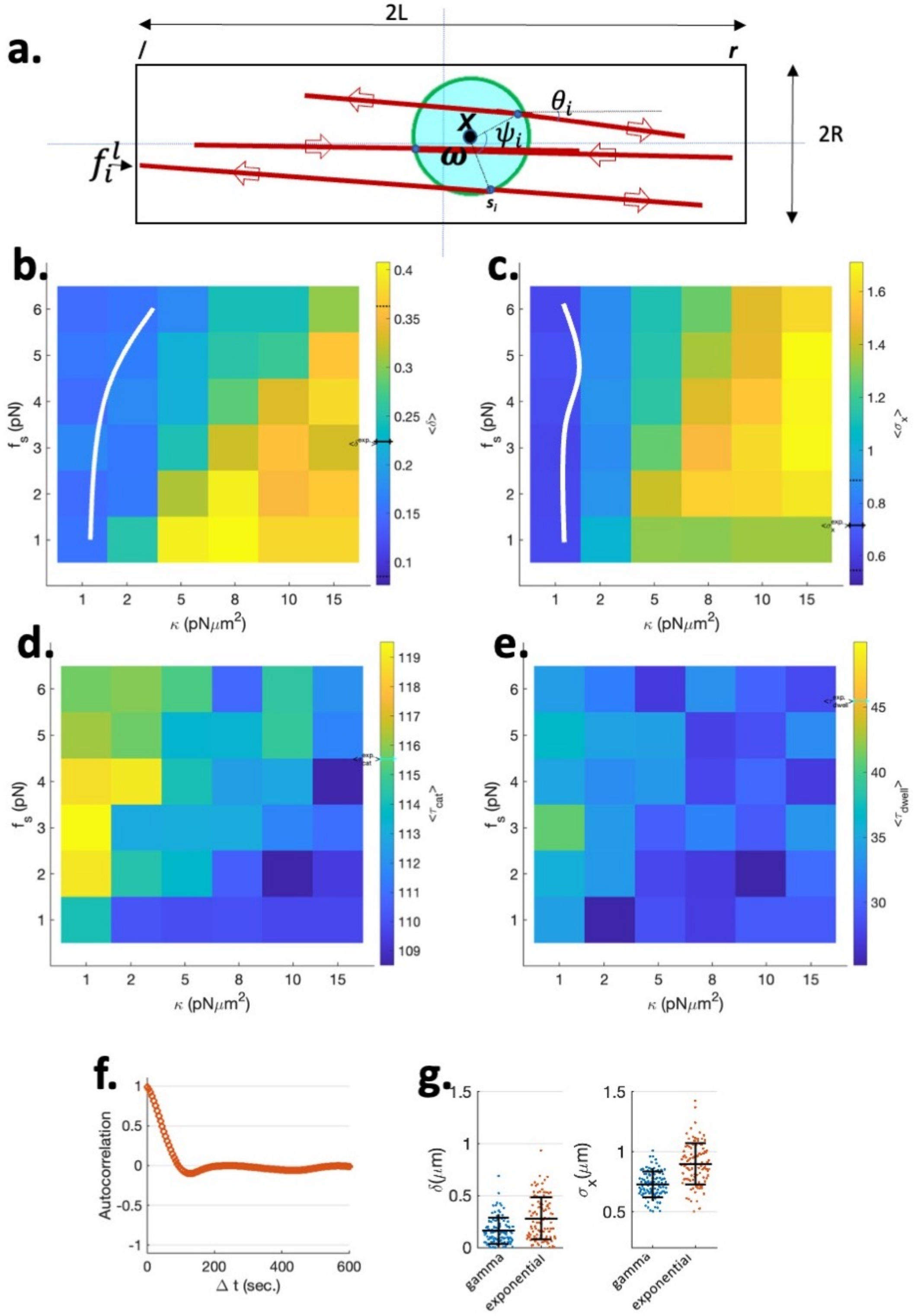


Figure S10: **a.** Schematic depicting elements of the theoretical model for MT-driven nucleus centering. The MTs originate from MTOCs (red dots) mounted on the periphery of a rigid nucleus. **b-c.** Discrete contour plots for $\langle\delta\rangle$ (b) and $\langle\sigma_x\rangle$(c) were obtained by systematically varying the only two unknown parameters in the model (average of 100 independent instances). The solid and dotted lines on the color bar respectively mark the experimental mean and standard deviation. The white counter line in (b) and (c) corresponds to the experimental mean value of $\delta$ and $\sigma_x$ respectively. **d.-e.** Discrete contour plots showing the mean catastrophe (d) and mean dwell time (e). The solid line on the color bar is the mean of experimental observation. **f.** Average auto-correlation from simulations using the parameter given in Fig 7c. **g.** Effect of shape of the $\tau_{cat}$ distribution on reliable and robust nuclear centering. Statistics of $\delta$ (left) and $\sigma_x$ (right) for the two catastrophe time distributions: gamma and exponential. The parameters in both cases are the same (as in Fig 7c) except for the $\tau_{cat}$ distribution parameters. The exponential distribution has the mean $\tau_{cat}$ same as the gamma distribution.
